# Disulfiram Protects Against Diet-Induced Obesity by Reprogramming Systemic Lipid Partitioning Independent of GSDMD

**DOI:** 10.64898/2026.02.06.704424

**Authors:** Babunageswararao Kanuri, Rohan R Varshney, Krishna P Maremanda, Nitin Nitin, Angelina Akomea, Dipanjan Chattopadhyay, Steve Ting-Yuan Yeh, Beatriz Y Hanaoka, Michael C Rudolph, Prabhakara R. Nagareddy

**Author notes:** **Corresponding author,** Dr. Prabha Nagareddy, PhD, FAHA, Department of Medicine, Cardiovascular Section, University of Oklahoma Health Sciences Center, Oklahoma City, USA Ex 44737.

## Abstract

Obesity remains a major global health challenge with limited durable pharmacotherapies. Disulfiram (DSF), an FDA-approved drug reported to inhibit gasdermin D (GSDMD), has been proposed to improve metabolic outcomes through suppression of inflammasome signaling. Here, we demonstrate that GSDMD is dispensable for high-fat diet–induced obesity and insulin resistance, as neither genetic deletion nor antisense-mediated inhibition of GSDMD confers metabolic protection. In contrast, DSF robustly protects against obesity and IR through a GSDMD-independent mechanism. These effects are not attributable to reduced caloric intake but instead reflect a coordinated reprogramming of systemic lipid handling. Under steady-state conditions, DSF suppresses basal lipid oxidation while promoting fecal fatty acid excretion. In striking contrast, during acute lipid challenge, DSF enhances tissue lipid utilization and accelerates systemic clearance. Together, these findings overturn the prevailing inflammasome-centric model and establish context-dependent regulation of lipid partitioning—rather than inflammasome inhibition—as the primary mechanism underlying DSF’s anti-obesity effects

## INTRODUCTION

Obesity is a chronic, relapsing, and progressive disease that has reached epidemic proportions in developed nations.^1^ It is now recognized as one of the most pressing health challenges of the modern era and a major contributor to cardiovascular disease (CVD), type 2 diabetes (T2D), and several forms of cancer.^2–4^ In the United States alone, more than 40% of adults and nearly 20% of children meet the clinical criteria for obesity.^5,6^ Lifestyle modification rarely produces durable weight loss, leaving many patients reliant on surgical or pharmacologic interventions that primarily target appetite suppression or caloric absorption. However, currently available FDA-approved anti-obesity medications remain limited, and their benefit–risk profiles often restrict long-term use.^7–11^ Given the rapidly rising prevalence of obesity and its substantial health and economic burden, there is an urgent need to develop safe, cost-effective, and sustainable therapeutic strategies. In this context, drug repurposing represents an attractive approach that accelerates clinical translation while minimizing safety concerns.^12,13^

Chronic low-grade inflammation within adipose tissue is a defining feature of obesity-associated insulin resistance (IR) and a major contributor to cardiometabolic disease.^14–16^ Crosstalk between innate immune cells and metabolic tissues sustains systemic inflammation and promotes excessive myelopoiesis.^17^ In particular, neutrophil-derived S100A8/A9 activates Toll-like receptor (TLR)-4 on adipose tissue macrophages, driving NLRP3 inflammasome activation and downstream interleukin-1β (IL-1β) signaling.^17^ Consistent with this framework, blockade of components of the NLRP3–IL-1β–IL-1R1 axis improves glycemic control and attenuates metabolic dysfunction in experimental models of obesity.^17–20^ Yet clinical translation has revealed important limitations: although IL-1β inhibition reduced major adverse cardiovascular events in the CANTOS trial, it failed to prevent progression to T2D in prediabetic individuals. ^21,22^ This discordance has been attributed to persistent activity of additional inflammatory mediators—including IL-18, TNF-α, IL-6, and other damage-associated factors—that continue to propagate metabolic dysfunction despite selective cytokine neutralization.^23^ These findings highlight the constraints of targeting individual inflammatory cytokines and underscore the need for strategies that more comprehensively modulate inflammasome-driven inflammatory programs.

Activation of the NLRP3 inflammasome culminates in pyroptosis, a lytic form of inflammatory cell death common to canonical and non-canonical inflammasome pathways. During this process, caspase-1 cleaves pro–IL-1β, pro–IL-18, and gasdermin D (GSDMD). The N-terminal fragment of GSDMD inserts into the plasma membrane and oligomerizes to form pores through which IL-1β, IL-18, and numerous intracellular inflammatory mediators are released. ^24–26^ Unlike strategies that neutralize individual cytokines downstream of inflammasome activation, inhibition of GSDMD targets the terminal execution step of pyroptosis and is therefore predicted to suppress the release of multiple inflammatory factors during both sub-lytic and lytic phases of cell death. ^27^ Recent unbiased high-throughput screening identified disulfiram (DSF) as a covalent inhibitor of GSDMD, blocking pore formation by modifying a critical cysteine residue required for pyroptotic membrane insertion. ^27^ We therefore hypothesized that targeting GSDMD—either genetically or pharmacologically with DSF—would confer metabolic protection through suppression of GSDMD-dependent inflammasome signaling.

Here, we demonstrate that this hypothesis is incorrect. Genetic deletion and antisense-mediated inhibition of GSDMD fail to protect against high-fat diet–induced obesity or IR. In contrast, DSF retains full metabolic efficacy in Gsdmd-deficient mice, establishing a GSDMD-independent mechanism of action. Mechanistically, DSF reprograms systemic lipid handling in a context-dependent manner: under steady-state conditions, it diverts dietary lipids toward fecal excretion, whereas during acute nutrient challenge it accelerates lipid utilization and clearance without altering caloric intake or basal energy expenditure. Collectively, these findings identify lipid partitioning—not inflammasome suppression—as the principal mechanism underlying DSF’s metabolic actions and uncover an unrecognized intestinal control point governing systemic lipid fate.

## RESULTS

### DSF protects against diet-induced obesity and IR

A predefined timeline was implemented to systematically evaluate obesity- and IR-related parameters across multiple study designs including DSF treatment and ASO-based *Gsdmd* inhibition in WT mice, as well as *Gsdmd* knockout (KO) models. These approaches enabled assessment of DSF-mediated effects as well as the metabolic consequences of *Gsdmd* inhibition or gene deletion per se (Figure S1A). To validate the metabolic effects of DSF, WT C57BL/6J mice were fed a 60% HFD for 16 weeks, a duration sufficient to induce obesity and capture early DSF-mediated weight effects (Figure 1A).^28^ Based on prior dose-response studies, DSF was administered at 200 mg/kg throughout the study.^28,29^ As expected, HFD feeding induced progressive BW gain beginning at 4 weeks, which was markedly attenuated in DSF-treated mice (HFDD) (Figure 1B). Reduced BW was accompanied by significant decreases in fat mass at both early (4 weeks; Figure S1B) and later (10 weeks; Figure 1C) time points, as well as smaller adipocyte size and a shift toward reduced adipocyte hypertrophy at study termination (Figure 1D).

**Figure 1.**
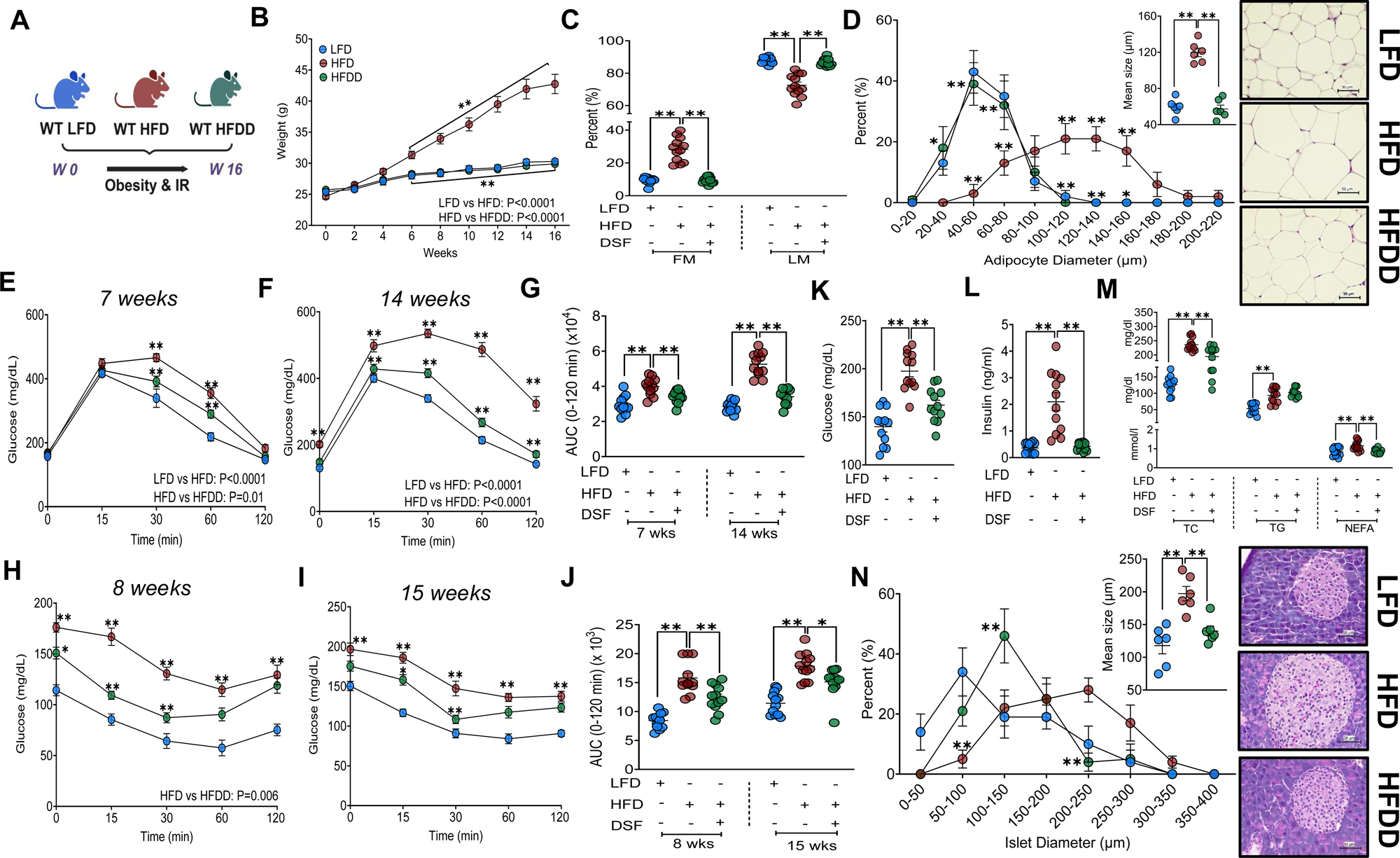
DSF protects against HFD-induced obesity and IR. **(A)** Experimental design for HFD feeding and DSF administration. **(B)** BW progression over 16 weeks in LFD, HFD, and HFDD mice. **(C)** Body composition at 10 weeks (FM and LM). **(D)** Representative H&E-stained adipose tissue sections with adipocyte size distribution and mean adipocyte area (n = 6/group). **(E-G)** GTTs performed at 7 and 14 weeks with corresponding AUCs. **(H-J)** ITTs performed at 8 and 15 weeks with corresponding AUCs. **(K-M)** Fasting blood glucose, plasma insulin, and circulating lipids (TC, TG, NEFA). **(N)** Representative pancreatic H&E sections with islet size distribution and mean islet area (n = 6/group). Data are mean ± SEM. *P < 0.05, **P < 0.01. n = 12 mice/group. Two-way ANOVA was used for BW, GTT, ITT, and adipocyte and islet size distribution. One-way ANOVA was used for all other comparisons unless stated. Kruskal-Wallis test was applied for fat mass, ITT (8-week AUC), and insulin.

DSF treatment also improved glucose homeostasis during obesity progression. HFD-fed mice exhibited impaired glucose tolerance (GTT) at both 7 and 14 weeks, whereas HFDD mice showed significantly improved glucose tolerance at both time points (Figures 1E-G). Consistent with this, insulin tolerance tests (ITT) performed at 8 and 15 weeks revealed enhanced insulin sensitivity in HFDD mice compared with HFD controls (Figures 1H-J). HFD feeding resulted in fasting hyperglycemia, hyperinsulinemia, and hyperglucagonemia, all of which were significantly normalized by DSF treatment (Figures 1K, 1L, and S1C). In parallel, HFD-induced elevations in circulating total cholesterol, triglycerides, non-esterified fatty acids (TC, TG, and NEFA) were significantly reduced by DSF treatment, indicating improved systemic lipid homeostasis (Figure 1M). Obesity-associated IR is characterized by compensatory pancreatic islet hypertrophy.^30^ Accordingly, increased circulating insulin levels in HFD mice were associated with enlarged pancreatic islets, a phenotype that was attenuated in HFDD mice (Figure 1N).

### DSF-induced metabolic protection is independent of dietary caloric intake

To determine whether reduced caloric intake contributes to DSF-mediated protection against obesity and IR, food intake was quantified using both manual measurements and metabolic cage-based feeding assessments, while energy balance (EB) was evaluated by indirect calorimetry (IDC) in WT C57 mice fed LFD, HFD, or HFDD diets. Manual intake was measured at diet initiation and after 32 weeks of chronic feeding, with the extended feeding period included to benchmark against prior IDC datasets.^28^ At diet initiation, caloric intake did not differ among groups at days 7 or 14 (Figure 2B), and after chronic feeding, HFD and HFDD mice similarly consumed comparable calories over a two-week period (Figure 2D). Despite similar caloric intake, HFDD mice exhibited minimal BW changes (0-5% by Day 14) at both early and chronic time points, whereas HFD mice showed modest BW gain during early diet exposure (Figures 2C and 2E). These findings were independently corroborated using metabolic cages (Figure 2F), which revealed comparable caloric and water intake and, consequently, similar EB between HFD and HFDD mice (Figures 2G-I). To further exclude a role for caloric restriction, pair-feeding studies were performed in an independent cohort (Figure S1D). Compared with HFD mice pair-fed to match HFDD intake, HFDD mice exhibited reduced BW and fat mass (Figures S1E and S1F) but did not show improvements in glucose tolerance (GTTs at 8 and 14 weeks; Figures S1G, S1H, and S1J). In contrast, HFDD mice demonstrated robust improvements in glucose disposal following one week of ad libitum feeding (Figures S1I and S1J), indicating that an acute nutrient stress under ad libitum conditions would induce IR in HFD but not HFDD mice. Collectively, these findings indicate that DSF-mediated protection against obesity and metabolic dysfunction is not attributable to reduced caloric intake, supporting a mechanism independent of dietary energy restriction.

**Figure 2.**
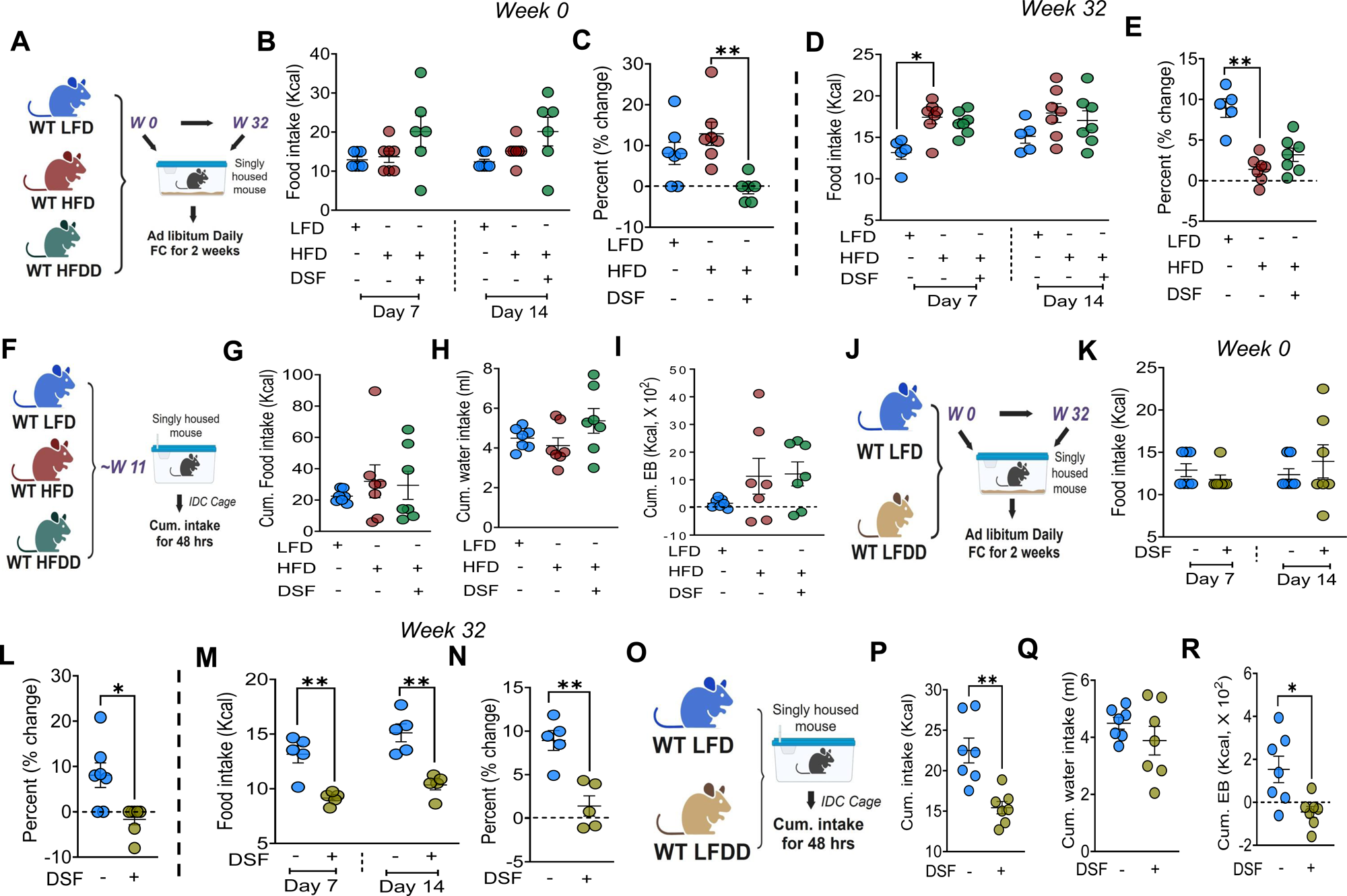
DSF-mediated protection against obesity is independent of caloric intake. **(A)** Schematic for manual food intake measurements in LFD-, HFD-, and HFDD-fed mice. **(B, C)** Caloric intake during diet initiation (days 7 and 14) and corresponding BW change. Kruskal–Wallis test for caloric intake. **(D, E)** Caloric intake and BW change following chronic (32-week) feeding. **(F-I)** Schematic and metabolic cage/IDC-based caloric intake, water intake, and EB measurements taken for 48 hrs ∼11 weeks of LFD, HFD, and HFDD dietary intervention. **(J-N)** Corresponding manual food intake experimental design, caloric intake, and BW change analyses performed in LFDD-fed mice. Mann–Whitney test for food intake and BW change, respectively, during diet initiation. n = 5/group for chronic feeding experiments. **(O-R)** Corresponding metabolic cage/IDC-based caloric intake, water intake, and EB measurements performed in LFDD-fed mice. Data are mean ± SEM. *P < 0.05, *P < 0.01. n = 6–7 mice/group unless otherwise stated. One-way ANOVA was used for multiple-group comparisons, and unpaired t test for two-group comparisons unless otherwise indicated.

To determine whether DSF per se affects caloric intake under low-fat dietary conditions, we also performed both manual and metabolic cage-based feeding assessments (Figures 2J and 2O). LFDD mice exhibited a significant reduction in food intake measured after 32 weeks of chronic feeding, but not during diet initiation (Figures 2K and 2M); this reduction was independently confirmed by metabolic cage-based measurements (Figure 2P). Consistent with decreased intake, LFDD mice displayed a negative EB relative to LFD controls (Figure 2R), although BW changes remained modest (0-5% by Day 14) during manual assessment periods (Figures 2L and 2N). Given the reduced caloric intake observed in LFDD mice, we next examined whether DSF directly modulates metabolic parameters under low-fat dietary conditions. Although LFDD mice displayed marked reductions in BW and fat mass, no significant differences were observed in glucose tolerance, insulin sensitivity, fasting glucose, circulating insulin levels, lipid profiles, or pancreatic islet morphology compared with LFD controls (Figures S2A-N). These data indicate that DSF per se does not exert overt metabolic effects under low-fat dietary conditions.

### DSF effectively inhibits pyroptosis-associated protein release in vitro

DSF has been reported to inhibit gasdermin D (GSDMD) pore formation,^27^ whereas small-molecule NLRP3 inhibitors such as MCC950 suppress upstream inflammasome activation.^31,32^ To directly compare the relative efficacy of these approaches in limiting pyroptosis-associated protein release, we performed unbiased proteomic profiling of the secretome from mouse bone marrow-derived macrophages (BMDMs) following inflammasome activation (Figure S3A). BMDMs were primed with LPS and treated with nigericin to induce pyroptosis, with DSF or MCC950 added after LPS priming. DSF dose optimization was performed using MTT-based viability assays and immunoblotting for GSDMD cleavage (Figures S3H-M, S3O). Higher DSF concentrations were cytotoxic (25, 50, and 100 µM), whereas lower doses failed to suppress pyroptosis (3.125 µM and 0.39 µM). A concentration of 10 µM preserved cell viability while effectively inhibiting GSDMD activation, as evidenced by reduced N-terminal cleavage, and was therefore used for all subsequent experiments. This dose matched the working concentration of MCC950 (see STAR Methods).

Proteomic profiling revealed robust alterations in the secretome following inflammasome activation. Compared with LPS alone (135 upregulated proteins; L vs DMSO), nigericin treatment significantly altered the abundance of hundreds of proteins, consistent with extensive pyroptotic cell death (234 upregulated and 618 downregulated; L+N vs L). MCC950 treatment broadly remodeled the secretome, reflecting widespread modulation of inflammasome-dependent signaling pathways (562 upregulated and 185 downregulated; L+MCC950+N vs L+N). In contrast, DSF treatment resulted in markedly fewer differentially abundant proteins, indicating a selective and effective inhibition of pyroptosis-associated protein release (66 upregulated and 4 downregulated; L+DSF+N vs L+N) (Figures S3B and S3C). Pathway enrichment analysis (LogFC ±1, adjusted P < 0.05) supported these findings. LPS treatment enriched pathways associated with inflammatory activation, including acute phase response and IL-1 signaling. Nigericin-induced pyroptosis further enriched pathways linked to cell death and loss of cellular homeostasis. MCC950 treatment produced broad pathway modulation, whereas DSF treatment resulted in minimal pathway enrichment, consistent with targeted suppression of pyroptotic membrane rupture rather than upstream inflammatory signaling (Figures S3D-G, S3N). Collectively, these data demonstrate that DSF selectively limits the release of intracellular proteins during pyroptosis in vitro, producing a more restricted alteration of the macrophage secretome compared with upstream inflammasome inhibition.

### GSDMD is dispensable for HFD-induced obesity and IR

Given that DSF inhibits GSDMD-mediated pyroptosis in vitro and protects against obesity in vivo, we next directly tested whether GSDMD itself contributes to diet-induced obesity and IR (Figure S4A). This was further motivated by prior reports demonstrating metabolic protection in *Nlrp3*-deficient mice under HFD conditions.^18,33^ Successful global deletion of *Gsdmd* was confirmed by near-complete loss of GSDMD protein in KO mice (Figure S4W). WT and KO mice were fed an HFD for 16 weeks. Contrary to expectations, KO mice exhibited body-weight trajectories (Figure S4B) indistinguishable from WT controls throughout the feeding period, with no differences in body fat mass (Figure S4C). Consistent with these findings, GSDMD deficiency did not improve glucose homeostasis, as assessed by GTTs (Figures S4D-F) and ITTs (Figures S4G-I) at two time points. Despite comparable adiposity, KO mice displayed significantly elevated fasting glucose (Figure S4J) and circulating insulin levels (Figure S4K), indicative of worsened glycemic control. Circulating TC, TG, and NEFA were similar between groups (Figure S4L). To determine whether altered energy homeostasis contributed to these findings, metabolic parameters were assessed by IDC (Figure S4M). WT and KO mice exhibited comparable caloric and water intake (Figure S4N), locomotor activity (Figure S4O), VO_₂_ and VCO_₂_ (Figure S4P), RER (Figure S4Q), EE (Figure S4R), and EB (Figure S4S). Manual food intake measurements during diet initiation similarly revealed no differences between groups (Figures S4T-V), indicating that neither early nor chronic alterations in energy intake or expenditure account for the lack of metabolic protection in *Gsdmd*-deficient mice.

Because lifelong gene deletion can elicit compensatory adaptations, we next employed ASO-mediated inhibition of *Gsdmd* to assess the effects of inducible gene suppression (Figure S6A). *Gsdmd* ASOs achieved robust, dose-dependent (15 and 30 mg/kg) knockdown of *Gsdmd* expression in liver, perigonadal white adipose tissue (pgWAT), and skeletal muscle (SKM) in chow mice, an effect that was maintained even in HFD mice (Figure S5B). Further, robust inhibition *of Gsdmd* occurred without inducing inflammatory toxicity or altering *Nlrp3* expression (Figures S5C and S5D). Despite effective gene suppression, *Gsdmd* ASO-treated mice fed an HFD exhibited increased BW (Figures S6B and S6C) and elevated fasting glucose (Figure S6K), with adiposity remaining comparable to controls (Figure S6D). Moreover, ASO treatment did not improve glucose tolerance (Figures S6E-G) or insulin sensitivity (Figures S6H-J). Metabolic cage studies (Figure S6L) further revealed reduced locomotor activity (Figure S6N) without changes in caloric intake, VO_₂_, VCO_₂_, RER, or EE (Figures S6M, S6O-Q). Together, these findings demonstrate that neither genetic deletion nor inducible inhibition of *Gsdmd* protects against HFD-induced obesity or IR, establishing that GSDMD is dispensable for diet-induced metabolic disease.

### GSDMD-independent metabolic benefits of DSF underscore its role in systemic metabolism

To determine whether the metabolic benefits of DSF require GSDMD, *Gsdmd* KO mice were fed an HFD with or without DSF supplementation (HFDD), and metabolic outcomes were compared across groups (Figure 3A). As expected, HFD-fed KO mice exhibited progressive BW gain beginning at 2 weeks and persisting through 16 weeks. In contrast, HFDD-fed KO mice were protected from weight gain and displayed significantly reduced body fat mass (Figures 3B and 3C). Histological analyses confirmed attenuation of adipocyte hypertrophy in HFDD-fed KO mice, with reduced mean adipocyte size relative to HFD-fed KO controls (Figure 3D). DSF treatment also markedly improved glucose homeostasis in HFD-fed *Gsdmd*-deficient mice. HFDD-fed KO mice demonstrated enhanced glucose tolerance at both 7 and 14 weeks (GTTs, Figures S6S, 3E, and 3F) and improved insulin sensitivity at 8 and 15 weeks (ITTs, Figures S6T, 3G, and 3H). These functional improvements were accompanied by significant reductions in fasting blood glucose and circulating insulin levels, along with decreased pancreatic islet size (Figures 3I-K).

**Figure 3.**
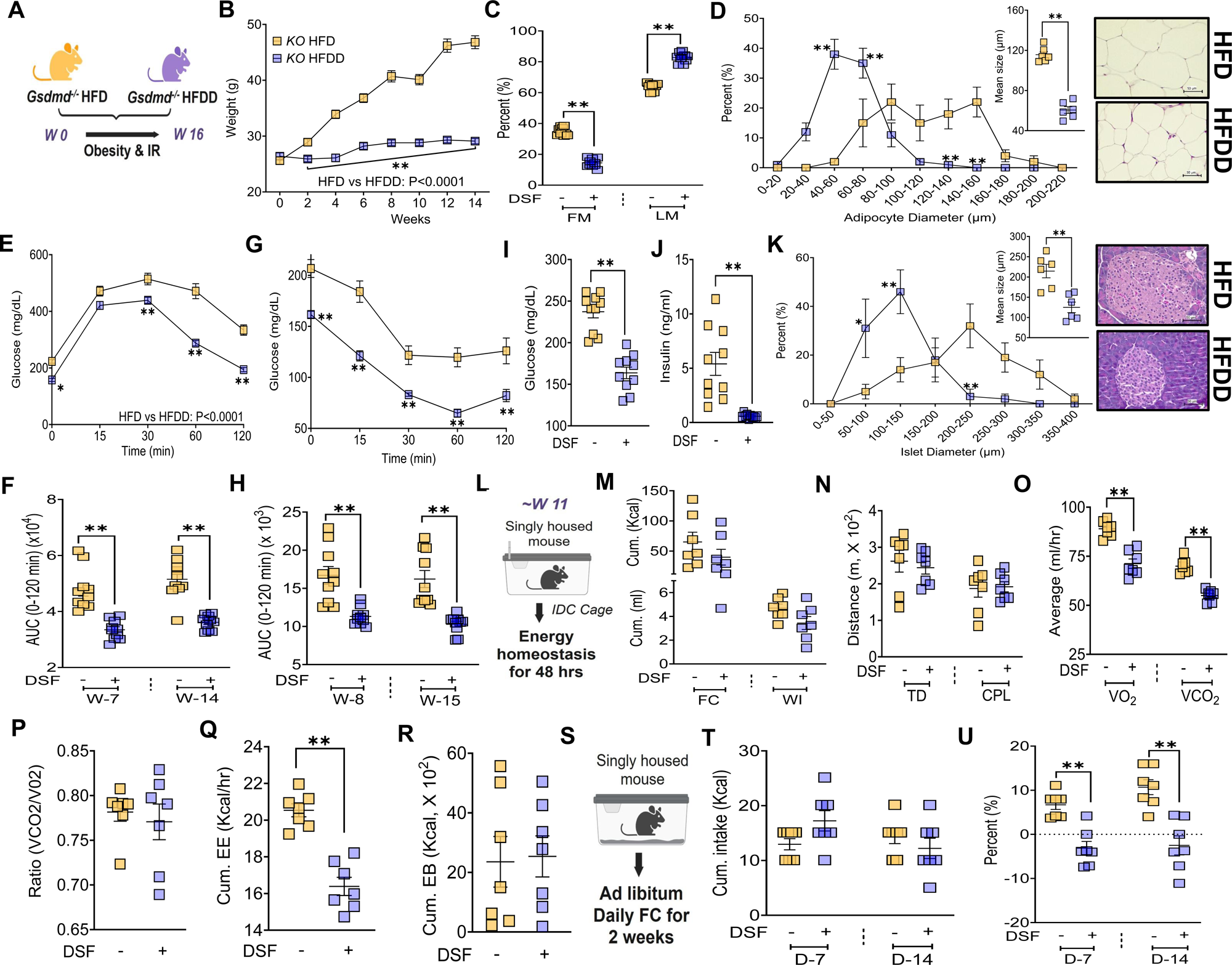
DSF-mediated metabolic protection is independent of GSDMD. **(A)** Experimental design for DSF treatment in *Gsdmd* KO mice. **(B)** BW progression during 16 weeks of HFD or HFDD feeding. **(C)** Body fat percentage at 10 weeks. **(D)** Adipose tissue H&E staining with adipocyte size distribution and mean adipocyte area (n = 6/group). **(E, F)** GTTs at 7 and 14 weeks with corresponding AUCs. **(G, H)** ITTs at 8 and 15 weeks with corresponding AUCs. **(I, J)** Fasting blood glucose and plasma insulin. **(K)** Pancreatic H&E staining with islet size distribution and mean islet area (n = 6/group). **(L-R)** Food intake, locomotor activity, and energy homeostasis parameters assessed by IDC (n = 7/group). **(S-U)** Manual food intake and BW change during diet initiation (n = 7/group). Data are mean ± SEM. *P < 0.05, *P < 0.01. n = 10 mice/group. Two-way ANOVA was used for BW, GTT, ITT, and adipocyte and islet size distribution; unpaired t test was used for two-group comparisons unless otherwise indicated. Mann-Whitney test was applied for ITT (15-week AUC), and fasting glucose.

To evaluate whether altered energy balance contributed to these effects, metabolic parameters were assessed by IDC (Figure 3L). HFDD- and HFD-fed KO mice exhibited comparable caloric and water intake (Figure 3M), locomotor activity (Figure 3N), RER (Figure 3P), and EB (Figure 3R). Manual food intake measurements during early dietary exposure further confirmed similar caloric consumption between groups (Figures 3S-U). Although VO_₂_, VCO_₂_, and overall EE were significantly reduced in HFDD-fed KO mice (Figures 3O and 3Q), these changes occurred in the absence of differences in energy intake or balance.

Collectively, these findings demonstrate that DSF protects against obesity and IR independently of GSDMD, indicating that its metabolic benefits are not mediated through GSDMD-dependent inflammasome pathways.

### DSF increases acute substrate utilization under lipid challenge

Despite comparable caloric intake and EB, WT mice fed HFD or HFDD differed markedly in lipid storage, prompting us to examine how ingested nutrients are processed under DSF treatment. To address this, we performed stable isotope tracer studies in WT mice fed experimental diets for ∼11 weeks, administering orally labeled lipid (¹³C_₁₆_-palmitate), carbohydrate (¹³C_₆_-glucose), or amino acid (¹³C_₆_-leucine) substrates (Figure 4A). HFDD mice exhibited significantly increased ¹³C exhalation following ¹³C_₁₆_-palmitate administration, reflected by an elevated 10-hr AUC, indicating enhanced fatty acid oxidation relative to HFD controls (Figures 4B and 4C). A similar increase in ¹³C exhalation AUC was observed following ¹³C_₆_-glucose administration (Figures 4F and 4G), whereas ¹³C_₆_-leucine oxidation was comparable between HFD and HFDD mice. In contrast, LFDD mice showed increased ¹³C exhalation only following ¹³C_₆_-leucine administration (Figures S7A and S7B). Analysis of tracer kinetics revealed altered rise and decay slopes for both ¹³C_₁₆_-palmitate (Figures 4D and 4E) and ¹³C_₆_-glucose (Figures 4H and 4I) selectively in HFDD mice, consistent with enhanced acute substrate utilization under DSF treatment. Although LFDD mice exhibited kinetic differences for ¹³C_₁₆_-palmitate, the absence of increased total AUC indicated minimal net enhancement of lipid utilization under low-fat dietary conditions. The markedly higher ¹³C exhalation observed in LFDD mice following ¹³C_₆_-leucine administration was similarly accompanied by significant differences in tracer kinetics (Figures S7C and S7D).

**Figure 4.**
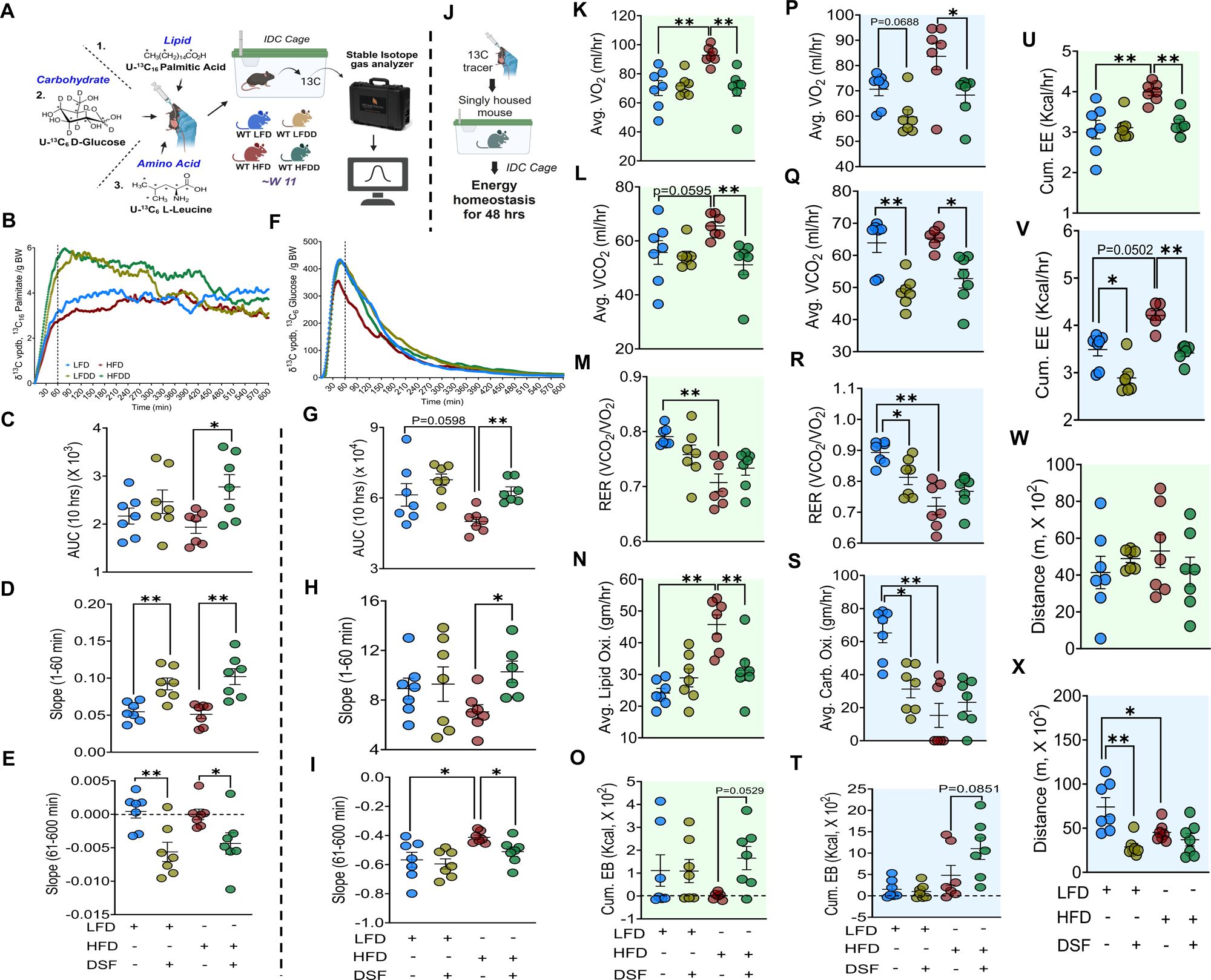
DSF enhances utilization of ingested substrates under acute challenge. **(A)** Schematic for oral administration of ¹³C-labeled tracers. **(B-E)** ¹³C exhalation following ¹³C_₁₆_-palmitate administration, corresponding AUC, and rise (1-60 min) and decay (61-600 min) slopes. **(F-I)** ¹³C exhalation kinetics following ¹³C_₆_-glucose administration with corresponding AUC and slope analyses. **(J)** IDC setup during tracer studies. **(K-M, O, U, W)** ¹²C-based VO_₂_, VCO_₂_, RER, cumulative EB, cumulative EE, and locomotor activity during ¹³C_₁₆_-palmitate administration. Kruskal-Wallis test for VCO_₂_ and EB. **(P-R, T, V, X)** ¹²C-based VO_₂_, VCO_₂_, RER, cumulative EB, cumulative EE, and locomotor activity during ¹³C_₆_-glucose administration. Kruskal-Wallis test for VO_₂_, VCO_₂_, cumulative EE, and cumulative EB. **(N, S)** Background lipid and carbohydrate oxidation indices. Kruskal-Wallis test for average carbohydrate oxidation during glucose. Data are mean ± SEM. *P < 0.05, *P < 0.01. n = 6–7 mice/group. One-way ANOVA was used unless otherwise stated.

To determine whether increased tracer oxidation reflected a generalized elevation in basal oxidative metabolism, we assessed background (¹²C-based) energy homeostasis during tracer studies using IDC (Figure 4J). HFDD mice consistently displayed reduced VO_₂_ (Figures 4K, 4P, and S7E), VCO_₂_ (Figures 4L, 4Q, and S7F), and cumulative EE (Figures 4U, 4V, and S7I) across substrates, while RER (Figures 4M, 4R, and S7G) and locomotor activity remained unchanged (Figures 4W, 4X, and S7J) relative to HFD controls. Basal lipid oxidation calculated in ¹³C_₁₆_-palmitate–administered groups was significantly reduced in HFDD mice (Figure 4N), whereas carbohydrate oxidation calculated in ¹³C_₆_-glucose administered groups remained unaffected (Figure 4S). EB was markedly increased in HFDD mice (Figures 4O, 4T, and S7H) despite lower EE. In contrast, DSF-treated LFD mice exhibited lipid oxidation comparable to LFD controls but significantly reduced carbohydrate oxidation (Figures 4N and 4S). EB in LFDD mice remained comparable to LFD controls across substrates (Figures 4O, 4T, and S7H), and changes in EE were substrate dependent (Figures 4U, 4V, and S7I). Moreover, the significant decrease in RER in LFDD mice suggested a shift in substrate preference (Figures 4R and S7G), a finding explored further under basal (non-tracer) conditions. Collectively, these findings demonstrate that DSF does not enhance basal lipid oxidation but selectively enhances oxidation of ingested substrates during acute nutrient challenge, supporting a context-dependent mechanism in which DSF promotes utilization of dietary lipids without increasing background metabolic demand.

### Tissue and plasma handling explain enhanced clearance under acute lipid challenge

The finding that DSF enhances lipid oxidation during acute lipid challenge, as revealed by ¹³C_₁₆_-palmitate tracing, raised three questions: whether unlabeled fatty acids in the vehicle (peanut oil) are similarly utilized, whether enhanced tracer oxidation reflects coordinated engagement of peripheral metabolic tissues, and whether a fraction of ingested lipids remains diverted toward excretion. These questions were addressed below.

Peanut oil, used as the vehicle for ¹³C_₁₆_-palmitate (5 mg/ml), contains a mixture of fatty acids (FA) such as saturated (Sat. FAs), monounsaturated (MUFAs), and polyunsaturated (PUFAs). Given the enhanced tracer utilization observed in HFDD mice, we hypothesized that these unlabeled fatty acids would also be preferentially cleared following DSF treatment. To test this, plasma lipidomic profiling was performed during the same 10-12 hr window after tracer administration (Figure 5A). Consistent with this hypothesis, HFDD mice exhibited significantly reduced circulating total FA (TFAs) compared with HFD controls (Figure 5B). Reductions were observed across multiple FA classes, including medium chain FA (MCFAs), Sat. FAs, MUFAs, and PUFAs (Figures 5C-F). Further stratification revealed significant decreases in long-chain, n-6, and n-3 PUFA subclasses (Figures 5G-I). The 16:1/16:0 ratio, a surrogate for SCD1 activity linked to de novo lipogenesis, was also significantly reduced in HFDD mice (Figure 5J). In contrast, DSF did not alter circulating total or individual FA species in LFD-fed mice. Together, these findings indicate that DSF selectively enhances circulating lipid clearance under HFD conditions.

**Figure 5.**
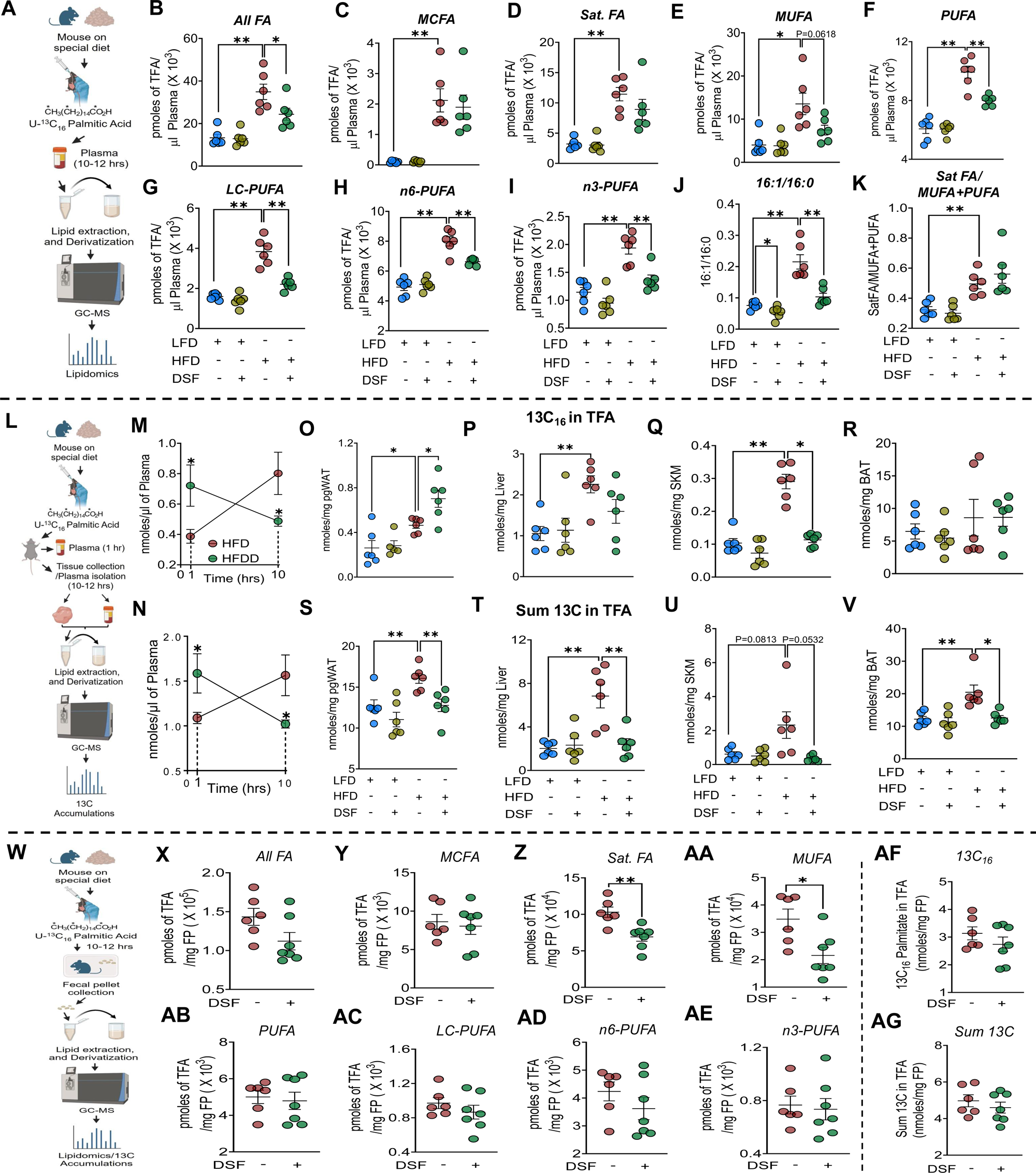
Acute lipid challenge promotes lipid utilization, but not excretion, in HFDD mice. **(A)** Experimental design for plasma lipidomic analysis following ¹³C_₁₆_-palmitate administration. **(B-F)** Plasma total and individual FA classes 10-12 hrs after lipid bolus. Kruskal–Wallis test for Sat. FA. **(G-I)** PUFA subclasses (long-chain, n-6, n-3). **(J, K)** Plasma lipid indices. **(L)** Experimental schematic for plasma and tissue ¹³C analysis. Mice fed LFD, LFDD, HFD, or HFDD were administered ¹³C_₁₆_-palmitate; blood was collected at 1 hr and at terminal time points (10-12 hrs), followed by collection of peripheral metabolic tissues. **(M, N)** Plasma ¹³C levels at 1 hr and terminal time points. Two-way ANOVA without multiple comparisons. **(O-R)** Tissue accumulation of ¹³C_₁₆_-palmitate in pgWAT, liver, SKM, and BAT. Kruskal–Wallis test for SKM. **(S-V)** Total ¹³C accumulation in corresponding tissues. Kruskal–Wallis test for BAT. **(W)** Experimental design for fecal lipidomics after ¹³C_₁₆_-palmitate administration. **(X-AE)** Fecal total and individual FA classes and PUFA subclasses. **(AF, AG)** Fecal ¹³C_₁₆_-palmitate and total ¹³C content. Data are mean ± SEM. *P < 0.05, *P < 0.01. n = 6–7 mice/group. One-way ANOVA was used for multiple-group comparisons, and unpaired t test for two-group comparisons unless otherwise indicated. The diet and drug description panels shown in Figures 5–7 apply to all panels within each corresponding column, including those without a top schematic.

To determine whether enhanced whole-body tracer utilization reflected tissue-specific handling, ¹³C enrichment was quantified in liver, perigonadal white adipose tissue (pgWAT), skeletal muscle (SKM), and brown adipose tissue (BAT) 10-12 hrs after tracer administration. Plasma ¹³C levels were additionally measured at both early (1 hr) and terminal time points to distinguish effects on uptake versus clearance (Figure 5L). Consistent with tracer kinetic analyses (Figures 4D and 4E), HFDD mice displayed higher plasma ¹³C_₁₆_-palmitate and total ¹³C at 1 hr post-administration (Figures 5M and 5N), indicating enhanced early appearance in circulation. In contrast, both measures were significantly reduced at the 10-12 hr time point consistent with accelerated systemic clearance.

Tissue-specific analysis revealed differential tracer pattern. HFDD mice showed a trend toward reduced ¹³C_₁₆_-palmitate in liver (Figure 5P), a significant reduction in SKM (Figure 5Q), and a significant increase in pgWAT (Figure 5O). To capture tracer-derived carbon flux beyond intact palmitate, total ¹³C was quantified. Notably, total ¹³C was markedly reduced across all tissues examined i.e., liver, pgWAT, SKM, and BAT in HFDD mice compared with HFD controls (Figures 5S–V), indicating enhanced intracellular utilization rather than prolonged retention. Supporting this interpretation, qPCR analyses performed 10-12 hr after tracer administration revealed increased expression of both lipid oxidation (*Ppar*α and *Pgc1*α) and lipogenesis (*Ppar*γ) markers selectively in pgWAT, but not in liver or SKM (Figures S7L-N).

We next assessed fecal FA composition and ¹³C enrichment in pellets collected 10-12 hr after tracer administration (Figure 5W). Analyses were restricted to HFD and HFDD groups, as LFD-fed DSF mice exhibited comparable plasma ¹³C and FA profiles as LFD mice (data not shown). Total fecal FA content did not differ between groups (Figure 5X); however, class-specific analysis revealed selective reductions in fecal Sat. FAs and MUFAs in HFDD mice (Figures 5Z and 5AA), with no changes in MCFAs or PUFAs, including their subclasses (Figures 5Y and 5AB-AE). Consistent with these findings, fecal ¹³C_₁₆_-palmitate and total ¹³C levels were similar between groups (Figures 5AF and 5AG), indicating that DSF does not enhance fecal lipid elimination during acute lipid challenge. Collectively, these data demonstrate that under acute lipid stress, DSF promotes rapid systemic lipid utilization and clearance through coordinated plasma and tissue handling rather than increased fecal elimination.

### Lipid excretion but not oxidation as a major driver of steady-state BW in HFDD mice

Basal lipid oxidation (¹²C-based) during lipid challenge remains unchanged despite marked enhancement of ¹³C_₁₆_-palmitate utilization, prompting us to determine whether lipid oxidation meaningfully contributes to the maintenance of steady-state BW in HFDD mice. To address this, we quantified whole-body substrate oxidation using IDCs performed ∼11 weeks of dietary intervention, in the absence of any acute substrate challenge (Figure 6A). Measurements were collected over cumulative 48-hr recordings and stratified by light and dark phases.

**Figure 6.**
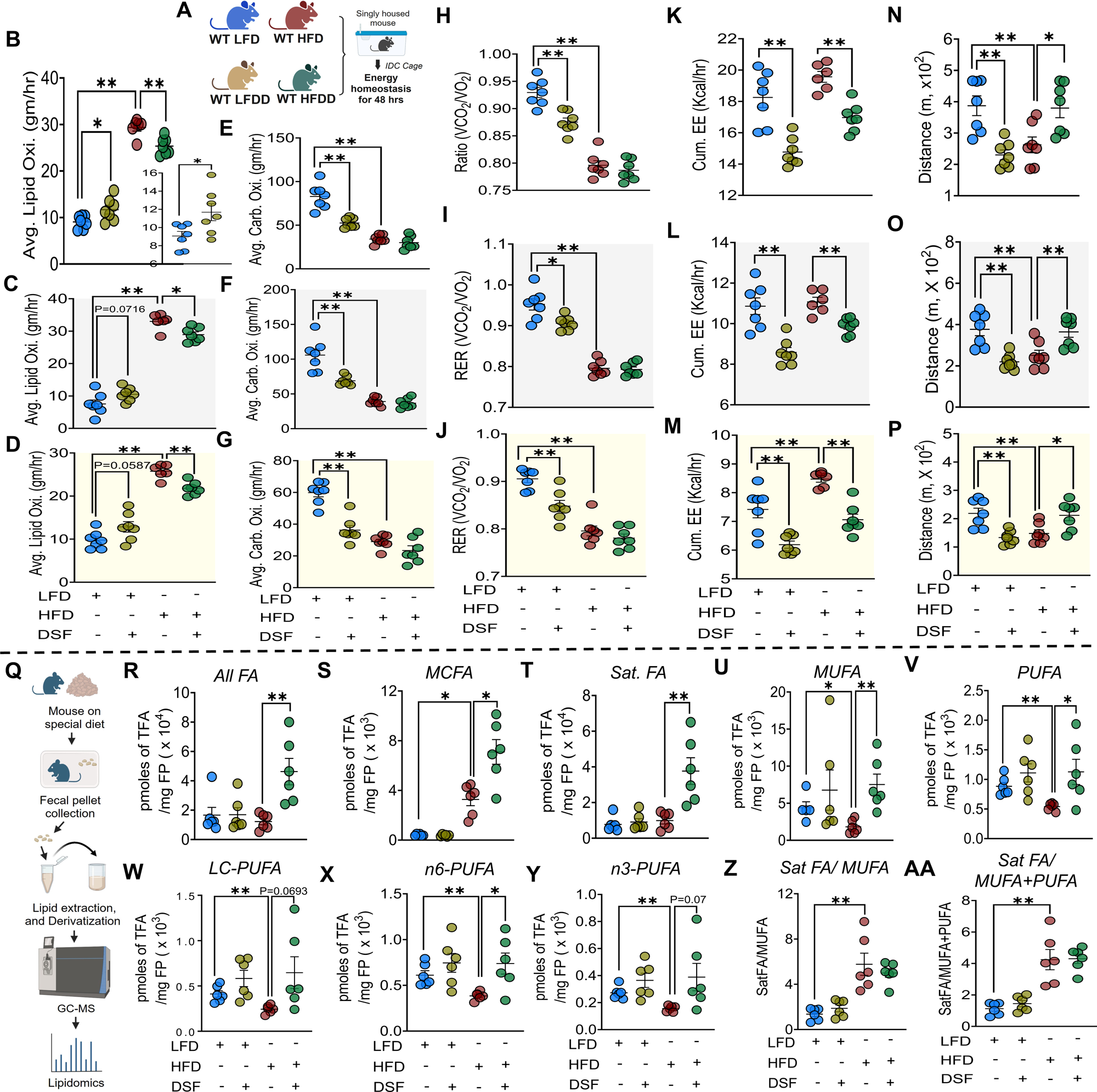
Basal lipid oxidation is reduced in HFDD mice and lipids are diverted to fecal excretion. **(A)** Experimental design for basal IDC measurements without tracer administration. **(B-D)** Lipid oxidation during cumulative, dark (active), and light (inactive) phases. **(E-G)** Carbohydrate oxidation across corresponding phases. **(H-J)** RER. **(K-M)** Cumulative EE. **(N-P)** Locomotor activity. **(Q)** Schematic for fecal lipidomic analysis. **(R-V)** Total and individual fecal FA classes, including MCFAs, Sat. FAs, MUFAs, and PUFAs. **(W-Y)** PUFA subclasses (long-chain, n-6, and n-3). Data are mean ± SEM. *P < 0.05, *P < 0.01. n = 6–7 mice/group. One-way ANOVA was used unless otherwise indicated. Kruskal-Wallis test was applied for TFA, Sat. FA, and MUFA; Mann-Whitney test was used for MCFA.

Contrary to expectations, DSF treatment significantly reduced lipid oxidation in HFD-fed mice across cumulative, light-phase, and dark-phase analyses (Figures 6B-D), while carbohydrate oxidation remained unchanged relative to HFD controls (Figures 6E-G). In contrast, LFDD mice exhibited increased lipid oxidation accompanied by reduced carbohydrate oxidation, indicating a diet-dependent divergence in basal substrate utilization (Figures 6B-G). DSF treatment reduced VO_₂_ (Figures S7O-Q), VCO_₂_ (Figures S7R-T), and EE (Figures 6K-M) regardless of dietary background, whereas RER was reduced only in LFDD mice (Figures 6H-J). Locomotor activity exhibited opposing diet-specific responses to DSF, with increased activity in HFDD mice and decreased activity in LFDD mice across both light and dark phases (Figures 6N-P). To dissociate metabolic effects from locomotion, metabolic parameters were analyzed during periods of minimal activity (see STAR Methods). Under resting conditions, DSF-treated mice continued to exhibit reduced VO_₂_, VCO_₂_, and EE independent of diet (Figures S7U, S7V, and S7X). RER remained unchanged between HFD and HFDD mice but was significantly reduced in LFDD mice relative to LFD controls (Figure S7W). Importantly, resting lipid and carbohydrate oxidation were both significantly reduced in HFDD mice, whereas LFDD mice exhibited a reduction only in carbohydrate oxidation (Figures S7Y and S7Z). Together, these findings demonstrate that basal lipid oxidation is reduced but not enhanced in HFDD mice, indicating that increased whole-body oxidative metabolism is unlikely to account for the maintenance of steady-state BW under DSF treatment.

Given these observations, we next explored alternative metabolic routes by which ingested lipids might be handled to preserve BW. In the context of comparable caloric intake and reduced whole-body oxidation, we hypothesized that enhanced fecal lipid excretion could serve as a compensatory mechanism. To test this, comprehensive lipidomic analyses were performed on fecal pellets collected ∼11 weeks of dietary intervention (Figure 6Q). Consistent with this hypothesis, DSF treatment in HFD-fed mice resulted in a marked increase in TFAs (Figure 6R). This increase was broad and non-selective, encompassing multiple FA classes, including MCFAs, Sat. FAs, MUFAs, and PUFAs (Figures 6S-V). Further stratification revealed significant increases in LC-PUFAs, n-6 PUFAs, and n-3 PUFAs (Figure 6W-Y). In contrast, DSF treatment in LFD-fed mice did not alter fecal excretion of total or individual FA species compared with LFD controls (Figures 6R-Y). Collectively, these findings identify fecal lipid excretion as a major metabolic route for disposal of ingested FAs in HFDD mice and highlight context-dependent remodeling of lipid partitioning as a key mechanism underlying DSF-mediated maintenance of stable BW.

To define the molecular basis underlying increased fecal lipid elimination under HFD conditions, qPCR-based gene expression analyses were performed in HFD-fed mice treated with DSF and compared with respective HFD controls, in the absence of any acute lipid challenge (Figures 7A). Expression of the lipid efflux transporters *Abcg5* and *Abcg8,* but not hepatic *Abcg1,* was significantly increased in both intestine and livers of HFDD mice (Figures 7B-C), implicating coordinated intestinal and hepatic pathways in DSF-induced lipid excretion. Further, marked induction of *Cyp2b10*, a diet-inducible enzyme that limits hepatic cholesterol accumulation via formation of polyhydoxylated bile acids, along with a trend toward increased *Cyp27a1* expression (p=0.0552), which catalyzes cholesterol conversion to bile acids (Figures 7D) was observed in livers of HFDD mice. Notably, these changes occurred without significant alterations in the expression of major regulators of both hepatic or intestine lipid uptake, oxidation, or clearance (Figures 7F-G) nor in genes encoding heparan sulfate proteoglycan-modifying enzymes (*Ndst1*, *Ext1*, and *Ext2*) (Figures 7E). Strikingly, enhanced lipid excretion was accompanied by reduced expression of key enzymes participating in de novo hepatic lipid synthesis, including *Fas* and *Hmgcr* (Figures 7H). Collectively, these data support a model in which DSF promotes cholesterol and fatty acid efflux with bile acid–mediated disposal while suppressing endogenous lipid synthesis, thereby maintaining steady-state BW under HFD conditions (Figures 7I).

**Figure 7.**
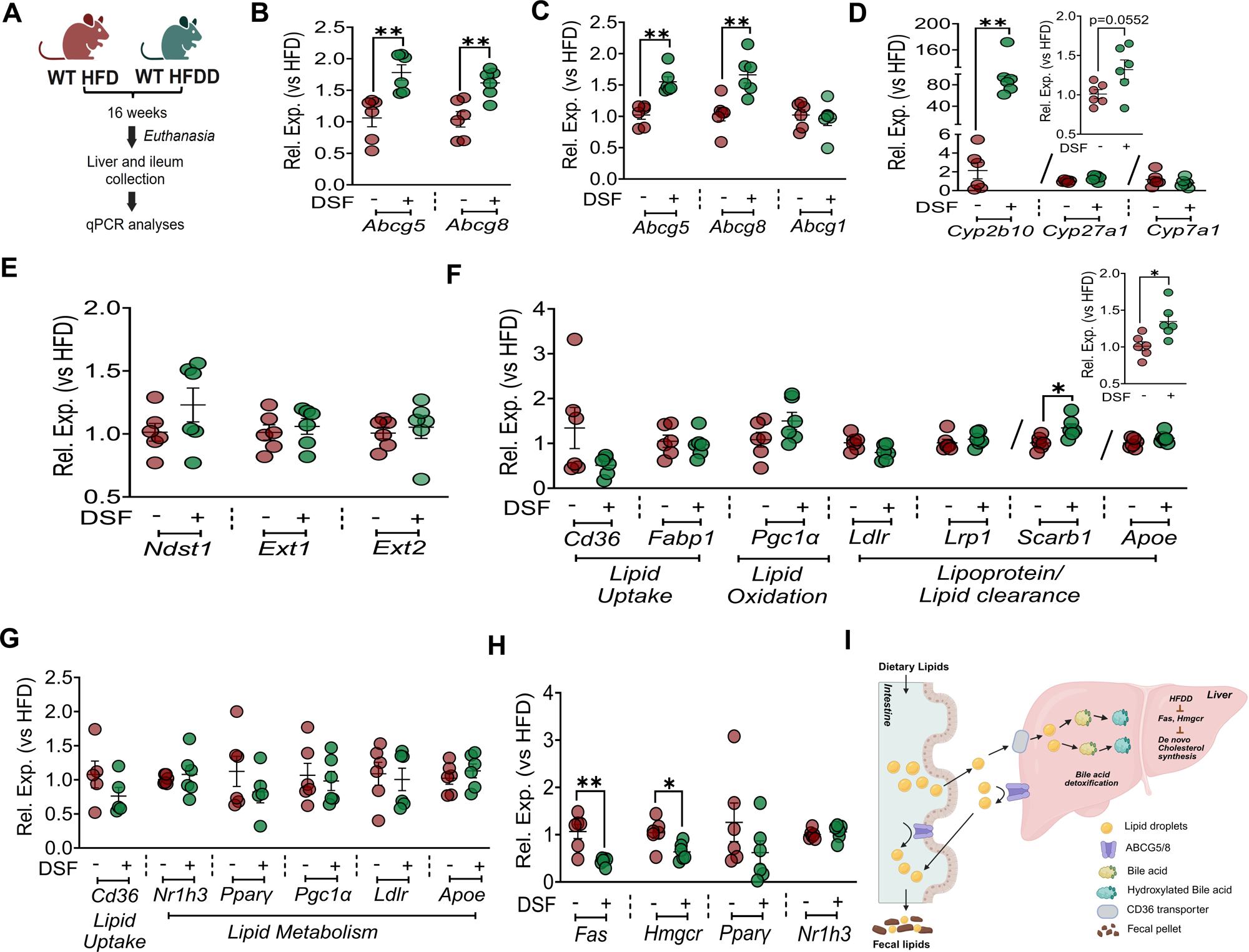
DSF promotes hepatobiliary and intestinal disposal of dietary lipids under HFD conditions. **(A)** Schematic for qPCR assessments of gene expressions. **(B, C)** Intestinal and hepatic expression of genes involved in lipid efflux. **(D, E)** Expression of genes regulating bile acid metabolism and enzymes modulating heparan sulfate proteoglycan (HSPG) function in liver. **(F, G)** qPCR analysis of genes involved in hepatic and intestinal lipid uptake, oxidation, and clearance. n=5 for intestinal *Cd36* and *Ppar*γ. **(H)** Hepatic expression of key lipogenic enzymes and transcriptional regulators. **(I)** Proposed model of DSF mediated lipid efflux with bile acid-mediated disposal while suppressing endogenous lipid synthesis, thereby maintaining steady-state BW under HFD conditions Data are mean ± SEM. *P < 0.05, **P < 0.01. n = 6 mice/group unless otherwise indicated. Unpaired t test was used unless noted; Mann-Whitney test for liver *Abcg5* and *Cyp2b10*.

## DISCUSSION

Our studies directly tested the prevailing hypothesis that suppressing adipose inflammasome execution through GSDMD inhibition would ameliorate diet-induced obesity and IR. Despite the well-established role of GSDMD in mediating pyroptotic pore formation and enabling the release of multiple pro-inflammatory cytokines, genetic deletion and antisense-mediated inhibition of GSDMD failed to confer metabolic protection in high-fat diet–fed mice. These findings indicate that blocking GSDMD-dependent inflammatory effector pathways is not sufficient to prevent obesity or IR in this setting. In contrast, we uncover a previously unrecognized, context-dependent metabolic mechanism by which DSF protects against diet-induced obesity and IR. We demonstrate that (i) GSDMD is dispensable for obesity and IR, (ii) DSF-mediated metabolic protection occurs independently of GSDMD, and (iii) DSF reprograms systemic lipid partitioning by diverting dietary fat toward fecal excretion under steady-state conditions while enhancing coordinated multi-organ lipid utilization during acute nutrient challenge.

Since the CANTOS trial, there has been renewed interest in identifying therapies that broadly mitigate cardiometabolic risk rather than selectively neutralize individual inflammatory mediators. ^23^Because DSF is FDA-approved, inexpensive, and well tolerated, it represents an attractive candidate for repurposing. Prior studies demonstrated that DSF protects against obesity and IR through mechanisms that remain incompletely defined^28,29,34^ and that it inhibits GSDMD pore formation in vitro,^27^. These observations fostered the prevailing assumption that its metabolic benefits are mediated through suppression of inflammasome signaling. However, our in vivo data directly challenge this interpretation. Neither global deletion nor ASO-mediated suppression of GSDMD protected against HFD-induced obesity or IR, and DSF retained full metabolic efficacy in Gsdmd-deficient mice. Together with recent reports dissociating GSDMD from obesity pathogenesis^35,36^ and evidence that DSF does not consistently suppress circulating IL-6 or TNF-α during chronic HFD feeding,^28^ these findings indicate that inflammasome signaling is unlikely to represent the principal mediator of DSF’s metabolic effects.

A central insight from our study is that DSF’s protective effects cannot be attributed to reduced caloric intake. Across manual measurements, metabolic-cage based assessments, and pair-feeding, HFD and HFDD mice maintained similar caloric intake. Importantly, earlier studies lacked pair-fed controls,^28,34^ leaving open the possibility that subtle differences in intake contributed to reduced BW gain^37–39^ and our findings rule out this explanation. Studies have shown that HFDD mice exhibited a marked reduction in FE,^28^ defined as BW gained per calorie consumed earlier,^37^ further revealing a dissociation between energy intake and fat storage. IDC analyses further clarified this dissociation. Despite reduced EE, HFDD mice maintained balanced EB, whereas LFDD mice exhibited negative EB. A balanced EB in HFDD mice indicates energy intake closely matched expenditure, whereas negative EB in LFDD mice reflects intake falling below expenditure.^40,41^ These diet-specific differences were accompanied by opposing changes in locomotor activity (both TD and PL, see STAR methods)^41^ and substrate oxidation. Under LFD conditions, DSF induced a caloric-restriction-like state characterized by reduced intake, decreased carbohydrate oxidation, and increased lipid oxidation. In contrast, under HFD conditions, DSF reduced basal lipid oxidation and EE while markedly increasing fecal lipid loss, indicating that enhanced lipid disposal but not oxidation that dominates steady-state BW maintenance in this context. Notably, decreased EE detected during states of enhanced locomotion as well as minimal activity dissociates EE with activity in DSF treated mice, which aligns with prior studies demonstrating that increased activity does not proportionally raise daily EE.^42^ At the molecular level, the coordinated upregulation of hepatic and intestinal efflux transporters (ABCG5/8), together with induction of regulators involved in bile acid detoxification, supports a model in which DSF enhances hepatobiliary and intestinal disposal of dietary lipids. Consistent with this framework, *Abcg5/8*-deficient mice exhibit impaired intestinal transport of dietary triacylglycerol and cholesterol into lymph,^43^ and HFD feeding protects against the development of fatty liver disease and loss of glycemic control independent of phytosterol accumulation.^44^ Concurrent suppression of endogenous lipid synthesis further constraints lipid accumulation, reinforcing a net negative lipid balance despite unchanged caloric intake.

A second, conceptually distinct insight from our work is that DSF selectively enhances utilization of ingested substrates during acute lipid challenge. Stable isotope tracer experiments revealed increased oxidation of orally administered ¹³C_₁₆_-palmitate and ¹³C_₆_-glucose without an accompanying rise in basal metabolic demand. Plasma lipidomics demonstrated rapid clearance of unlabeled dietary FAs present, confirming that DSF broadly enhances systemic lipid utilization rather than selectively metabolizing tracer substrates. Tissue ¹³C profiling further revealed coordinated tracer handling across liver, SKM, pgWAT, and BAT, characterized by reduced retention and accelerated turnover of tracer-derived carbon. Although we quantified “Sum 13C” to capture tracer-derived carbon flux beyond intact palmitate, recognizing that 13C generated via β-oxidation may enter other lipid or intermediary metabolic pathways,^45^ comprehensive mapping of these downstream fates remains an important direction for future studies. Importantly, fecal ¹³C levels were unchanged under acute challenge, indicating that enhanced oxidation, rather than excretion, predominates under conditions of lipid excess. EB during tracer administration remained dynamically positive in HFDD mice, consistent with substrate availability exceeding oxidative capacity within the 10-12 hr post-administration window. Moreover, VO_2_, VCO_2_, RER, and EE remained largely unchanged during tracer exposure, indicating that DSF’s effects on basal metabolism persist even during periods of acute nutrient surplus. Together with fecal lipidomic data obtained under steady-state conditions, these results support a dual-mode model of DSF action: under basal conditions, dietary lipids are preferentially diverted toward excretion, whereas during acute lipid surges, they are routed toward oxidation and clearance. This context-dependent regulation of lipid partitioning provides a unifying explanation for how DSF limits fat accumulation without reducing caloric intake or increasing basal EE, a mechanism that may be particularly relevant during episodes of overconsumption, such as post-fasting refeeding.^46^

In conclusion, our findings establish DSF as a metabolic reprogramming agent that redirects lipid fate rather than suppressing appetite or inflammation. By decoupling energy intake from fat storage, DSF identifies lipid partitioning as a druggable axis for obesity therapy and redefines its mechanism of action as independent of inflammasome signaling.

While this study provides mechanistic insight into DSF-mediated protection against obesity and IR, several limitations should be acknowledged. Although ¹³C_₁₆_-palmitate tracer experiments demonstrate enhanced lipid utilization under acute substrate challenge, they do not fully resolve the downstream metabolic fate of tracer-derived carbon, which will require comprehensive fluxomic and pathway-resolved analyses to define specific metabolic routing. In addition, while fecal lipidomic profiling indicates increased FA excretion under steady-state conditions, the precise intestinal and hepatic mechanisms responsible for this lipid diversion were not directly interrogated. Whether DSF acts primarily through modulation of sterol transporters, bile acid synthesis, chylomicron assembly, or coordinated enterohepatic signaling remains to be established through targeted genetic and mechanistic studies. A further limitation is that our findings do not fully reconcile the apparent disconnect between upstream and downstream inflammasome components in metabolic disease. Genetic deletion of Nlrp3 has been reported to confer protection against diet-induced obesity and IR in multiple models^18^, whereas our data demonstrate that deletion or inhibition of the downstream executioner GSDMD does not. The reasons for this divergence remain unclear. It is possible that NLRP3 influences metabolic homeostasis through non-canonical functions independent of pyroptotic pore formation, or through inflammasome-independent scaffolding or transcriptional effects. Alternatively, compensatory pathways downstream of caspase activation may preserve inflammatory signaling in the absence of GSDMD. Resolving this mechanistic discrepancy will require future studies dissecting GSDMD-independent outputs of NLRP3 signaling in metabolic tissues. Finally, indirect calorimetry (IDC)-based assessments rely on steady-state assumptions; therefore, interpretation of oxidation parameters during acute tracer administration should be approached with caution.

## Supporting information

Reagents and Resources

## RESOURCE AVAILABILITY

### Lead contacts

Further information and requests for resources should be directed to and will be fulfilled by the lead contacts, Babunageswararao Kanuri and Prabhakara Nagareddy.

### Materials availability

*Gsdmd* ASO used in the current study was provided by IONIS pharmaceutics under a material transfer agreement. All other materials used in this study that are commercially available are specified in the key resources table.

### Data and code availability

All data that support the findings of this study are available within the paper and its supplemental information.

## ACKNOWLEDGMENTS

This work was supported by funds from the NIH (HL137799, HL156856) and AHA (TPA97002) to PN. Histology service provided by the Stephenson Cancer Center tissue pathology core was supported by the National Institute of General Medical Sciences Grant P30GM154635 and National Cancer Institute Grant P30CA225520 of the National Institutes of Health”. The experimental schematics provided in different figures were created with BioRender.com.

## AUTHOR CONTRIBUTIONS

Conceptualization and research design, B.N.R.K., and P.R.N., methodology, B.N.R.K., R.R.V., M.C.R., S.T-Y.Y. and P.R.N., investigation and formal analysis - B.N.R.K., R.R.V., K.P.M., N.N., A.A., D.C., writing - original draft, B.N.R.K., and P.R.N., writing - edit draft, B.N.R.K., K.P.M., B.Y.H., and P.R.N., resources, S.T-Y.Y., M.C.R., and P.R.N. funding acquisition and supervision, P.R.N.

## DECLARATION OF INTERESTS

None

## Notes

### Competing Interest Statement

The authors have declared no competing interest.

