## Supplementary material for "Disulfiram Protects Against Diet-Induced Obesity by Reprogramming Systemic Lipid Partitioning Independent of GSDMD": Methods, Online Figure Legends

#### Animals, Husbandry, Diets, and Dietary Intervention

Procedures involving animal experimentation were approved by Institutional Animal Care and Use Committees (IACUCs) at the Ohio State University (OSU) as well as University of Oklahoma Health Sciences Center (OUHSC), in accordance with the NIH Guide for the Care and Use of Laboratory Animals. 8-12 weeks old wildtype male C57BL/6J mice and *Gasdermin D* knockout (KO) mice were purchased from Jackson Laboratories (Bar Harbor, MN, USA). All mice were housed in IVC cages, maintained at standard housing conditions and provided *ad libitum* access to food and water. Customized high-fat diets, with (HFDD; 5.03 kCal/g), or without drug incorporation (HFD; 5.04 kCal/g), along with corresponding low fat control diets, with (LFDD; 3.75 kCal/g) or without (LFD; 3.76 kCal/g) drug supplementation were procured from Dyets Inc. Previous studies have maintained HFD feeding in mice for up to 64 weeks, despite measurable decrease in BW due to DSF treatment occurring within the initial weeks of dietary intervention. [1]Accordingly, we selected a 16-week feeding period as an optimal timeframe to assess DSF-mediated effects on obesity development. Reported dose comparison studies revealed that DSF administered at 200 mg/kg produced more pronounced metabolic benefits than a 100 mg/kg dose under HFD conditions;[1,2] therefore, the higher dose (200 mg/kg; HFDD) was used for all experiments performed here. Fasting was performed for biochemical assessments, while tissue collection following euthanasia for molecular biology (qPCR/Western blots) was performed under normal feeding cycle (i.e., *ad libitum*) without any fasting.

In case of antisense oligonucleotides (ASO) based gene inhibition studies, C16 Palmitate-conjugated ASO targeting mouse Gasdermin D was developed and provided by Ionis Pharmaceuticals. Initial dose characterization studies for *Gsdmd* ASO were performed using two arbitrary doses (15 and 30 mg/kg), selected empirically with consideration of the extended half-life and altered tissue partitioning conferred by C16 (palmitic acid) conjugation and the expanded adipose compartment in obesity.[3,4] Mice (N=3) received single ASO injection via s.c. route on D0, D5 and D-10, respectively (Figure S4A). Control (CTL) ASO group received a single dose of 30 mg/kg. Tissues were collected from euthanized mice on D-14 for qPCR expression analyses of *Nlrp3* and *Gsdmd* genes. Tissue *Mcp1* expression was also quantified as an inflammatory biomarker to monitor potential C16 ASO-induced toxicity.[5] For actual experiments involving HFD feeding, both *Gsdmd* and CTL ASOs were injected sub-cutaneous (s.c.) once a week for 16 weeks.

For paired-feeding experiments, HFD mice were provided with the same amount of food consumed daily by HFDD mice for a total duration of 14 weeks. Briefly, HFDD mice were singly housed, and their daily food intake (g) was recorded throughout the study. Based on these measurements, an equivalent amount of food was randomly assigned and provided to singly housed HFD mice during the corresponding days of the subsequent week. Food allotments for HFD mice during week 0 (W 0) were determined using food intake measurements from each individual HFDD mouse recorded during the week preceding the start of the actual experiment (W-1). Upon completion of the 14-week paired-feeding period, all mice were transitioned to an *ad libitum* feeding regimen for an additional week (W 15).

Dietary intervention was also performed in a small cohort of mice that had bone marrow transplantation (BMT) as described previously.[6] Briefly, following BMT, mice were allowed to reconstitute for ~ 5 weeks, after which they were subjected to an 8-week HFD regime.

**Body composition analysis**

Body composition was assessed in different mouse groups as a primary measure of phenotypic characterization for HFD-induced obesity using quantitative magnetic resonance (qMR; EchoMRI® Whole Body Composition Analyzer 4-in-1/500, Echo Medical Systems, Houston, TX,

USA). During scanning, radio-frequency pulses were applied at a defined static magnetic field, and mice were allowed limited vertical and horizontal movement throughout the measurement period.[7]

**Indirect calorimetry and Stable isotope gas analysis**

Mice were placed in indirect calorimetry chambers (IDC, Sable Systems International, Las Vegas, Nevada) to assess the changes in multiple metabolic parameters such as calorie intake, $VO_2$ and $VCO_2$, respiratory exchange ratio (RER, $VCO_2/VO_2$), locomotion, and energy balance (EB). Data was analyzed using CalR, a web-based analysis tool for indirect calorimetry experiments (calrapp.org).[8] Indices such as carbohydrate oxidation (gm/hr), lipid oxidation (gm/hr), and energy expenditure (EE, kCal/hr) were calculated using $4.585 \times VCO_2 - 3.226 \times VO_2$, $1.695 \times VO_2 - 1.701 \times VCO_2$, and $0.06 \times (3.941 \times VO_2 + 1.106 \times VCO_2)$, respectively.[7] For locomotor activity data analyses, both total distance (TD) that includes X, Y and Z axes movement, and pedestrian locomotion (PL) that excludes Z axes movement from others were considered. To determine the metabolic changes during resting states, we selected five distinct 1 hr time points characterized by negligible locomotor activity during light phase of the day to analyze corresponding metabolic readouts.

In separate experiments, mice placed in IDC chambers were gavaged with 100 mg/kg BW of uniformly labeled Palmitic acid (U-$^{13}C_{16}$, 5 mg/ml dissolved in peanut oil)[9] or uniformly labeled Glucose (U-$^{13}C_6$, 1 g/kg BW in sterile water)[10] or uniformly labeled Leucine (U-$^{13}C_6$, 200 mg/kg BW in sterile water)[11] based on earlier reports. In addition to $13C_{16}$-palmitic acid, $13C_6$-glucose and $13C_6$-leucine were selected because obesity and IR are associated with altered glucose and amino acid metabolism. The labeled ($^{13}C$) $CO_2$ exhaled by mice per minute was measured using a stable isotope analyzer (Sable Systems International). This provided a measure of palmitate, leucine and glucose oxidation by the mice. For $^{13}C$ tissue accumulation studies, mice gavaged with 100 mg/kg BW U-$^{13}C_{16}$-Palmitic acid were euthanized 10-12 hrs post administration and multiple tissues (liver, skeletal muscle, perigonadal white adipose tissue, ands brown adipose tissue) were collected to measure $^{13}C_{16}$-palmitate utilization by GC-MS.

**Gas chromatography - mass spectrometry (GC-MS) analyses**

Tissues (10-100 mg) were added to 2 ml bead mill tubes containing 500 ul of 100 mM potassium phosphate buffer pH 6.8 and 500 ul methanol, and exact tissue masses were recorded.[12,13] Following bead mill homogenization, the supernatant was transferred to 13x100 mm borosilicate glass tubes. Homogenates were acidified with 40ul of prewashed 1N HCl (with hexanes) and vortexed briefly. Total lipids were extracted twice by adding 1ml of isooctane:ethyl acetate 3:1 (vol/vol) and once with 1ml Hexanes; for each extraction, samples were vortexed vigorously for 10-15 sec and centrifuged at 2000 g for 1 min to complete phase separation. The upper organic layer from each extraction was combined into a new 13x100 mm borosilicate glass tube. Extracted lipids were brought to dryness and resuspend in 300 to 1000 µl 2,2,4-Trimethylpentane (isooctane). For the unlabeled non-esterified fatty acid (NEFA) fraction, 20 to 100 µl of the resuspended total lipids was transferred to a new borosilicate glass tube, mixed with (25 ng) blended stable isotope internal standard, taken to dryness under gaseous $N_2$, and resuspended in 25 µl of 1% pentafluorobenzyl bromide in acetonitrile to which 25 µl of 1% diisopropylethylamine in acetonitrile was added and samples were incubated at room temperature for 30 minutes. Pentafluorbenzyl-fatty acid derivatives were taken to dryness under gaseous $N_2$ and resuspended in 100 µl hexanes. For the unlabeled total fatty acid fraction (TFA), 20 to 50 µl of the original total lipid extract was transferred to a separate Teflon lined screw cap glass tube, mixed with (75 ng) blended stable isotope internal standard, and taken to dryness under gaseous $N_2$. TFA samples were resuspended in 500 µl of 100% ethanol to which 500 µl of 1M NaOH was added to saponify the TFA fraction at 90°C for 30 min, followed by acidification using 550 µl of 1M HCl. Saponified samples were then extracted twice with 1.5 ml of Hexanes, taken to dryness under gaseous $N_2$, and derivatized as above. Derivatized TFA samples were resuspended in 300 µl hexanes for injection.

For $13C_2$ acetate tracer incorporation analysis, 20 to 50 µl of the resuspended total lipids was transferred to a new Teflon lined screw cap glass tube, mixed with (75 ng) d31 palmitate internal standard and taken to dryness under gaseous $N_2$. The samples were saponified, extracted and derivatized as above. Derivatized samples were resuspended in 300 µl hexanes for injection. For the NEFA, TFA and tracer fractions, 1 µl of pentafluorbenzyl-fatty acid derivatives was injected and data were collected on the GC-MS (8890 GC, 5977B MSD, Agilent) DB-1MS UI column (122-0112UI, Agilent) with the following run program: 80°C hold for 3 min, 30°C/min ramp to 125°C, no hold, 25°C ramp to 320°C and hold for 2 min. The flow for the methane carrier gas was set at 1.5 ml/min. Data was acquired in full scan negative chemical ionization mode to identify fatty acids of acyl chain length from 8 to 22 carbons. Peak areas of the analyte or of the standard were measured, and the ratio of the area from the analyte-derived ion to that from the internal standard was calculated. Tracer ($13C_2$ acetate) incorporation in palmitic acid was analyzed in two carbon increments from $13C_2$ to $13C_{16}$ against the d31-palmitate internal standard.

**Dose optimization studies for secretome proteomics analysis**

A working concentration of 10 µM MCC950 was selected based on prior studies from our laboratory demonstrating optimal pharmacological efficacy at this dose. For DSF, dose optimization was performed using MTT-based cell viability assays and western blot analysis (Figure S3H). BMDMs were treated with serial dilutions of DSF (100 µM to 0.39 µM; Figure S3I) under two conditions: DSF added prior to LPS priming (Figure S3J) or DSF added after LPS incubation (Figure S3L).

Across both conditions, DSF exhibited dose-dependent cytotoxicity, with higher concentrations (25, 50, and 100 μM) consistently reducing cell viability and therefore considered lethal (Figures S3K and S3M). Prolonged DSF exposure (30 min pre-LPS incubation) reduced viability even at lower doses, with cell survival averaging ~75% (Figure S3K), whereas shorter exposure (DSF added after 3 hrs of LPS priming) preserved viability up to ~95% at the lowest concentrations tested (Figure S3M). Notably, the inhibitory effects of DSF on pyroptotic cell death were significantly diminished at lower concentrations in the presence of LPS and nigericin, irrespective of whether DSF was added before or after LPS priming (Figures S3K and S3M). Based on these findings, DSF doses between 3.125 μM and 0.39 μM were deemed suboptimal. A working concentration of 10 μM was selected because it (i) matched the dose used for MCC950 and (ii) fell within the range (6.25–12.5 μM) that maintained optimal cell viability under both treatment conditions (Figures S3K and S3M). This dose was further validated by western blot analysis, which demonstrated that 10 μM DSF effectively inhibited GSDMD activation, as evidenced by reduced N-terminal cleavage. As expected, LPS and nigericin (L+N) treatment induced robust GSDMD cleavage, reflected by reduced levels of full-length GSDMD and the appearance of the cleaved fragment (Figure S3O). Based on these results, subsequent secretome analyses were performed with both DSF and MCC950 added after 3 hrs of LPS priming at a concentration of 10 μM (Figure S3A).

**Secretome proteomics on Orbitrap Exploris DIA - 12 Th windows**

Total protein from each sample was reduced, alkylated, and purified by chloroform/methanol extraction prior to digestion with sequencing grade modified porcine trypsin (Promega).[14] Tryptic peptides were trapped and eluted on 3.5 μm CSH C18 resin (Waters) (4 mm x 75 μm) then separated by reverse phase X Select CSH C18 2.5 um resin (Waters) on an in-line 150 x 0.075 mm column using an UltiMate 3000 RSLCnano system (Thermo). Peptides were eluted at a flow rate of 0.300 µL/min using a 60 min gradient from 98% Buffer A (0.1% formic acid, 0.5% acetonitrile) : 2% Buffer B (0.1% formic acid, 99.9% acetonitrile) to 95:5 at 2.0 min to 80:20 at 39.0 min to 60:40 at 48.0 min to 10:90 at 49.0 min and hold until 53.0 min and then equilibrated back to 98:2 at 53.1 min until 60 min. Eluted peptides were ionized by electrospray (2.4 kV) through a heated capillary (275°C) followed by data collection on an Orbitrap Exploris 480 mass spectrometer (Thermo Scientific). Precursor spectra were acquired with a scan from 385-1015 Th at a resolution set to 60,000 with 100% AGC, max time of 50 msec, and an RF parameter at 40%. DIA was configured on the Orbitrap 480 to acquire 50 x 12 Th isolation windows at 15,000 resolution, normalized AGC target 500%, maximum injection time 40 ms). A second DIA was acquired in a staggered window (12 Th) pattern with optimized window placements.

Following data acquisition, data were searched using Spectronaut (Biognosys version 19.1) against the UniProt *Mus musculus* database (Proteome ID: UP000000589, Taxon ID: 10090, 4$^{th}$ version of 2024) using the directDIA method with an identification precursor and protein q-value cutoff of 1%, generate decoys set to true, the protein inference workflow set to maxLFQ, inference algorithm set to IDPicker, quantity level set to MS2, cross-run normalization set to false, and the protein grouping quantification set to median peptide and precursor quantity.[15] Fixed Modifications were set to  Carbamidomethyl (C) and variable modifications were set to Acetyl (Protein N-term), Oxidation (M). Protein MS2 intensity values were assessed for quality using ProteiNorm.[16] The data was normalized using VSN[17] and analyzed using proteoDA to perform statistical analysis using Linear Models for Microarray Data (limma) with empirical Bayes (eBayes) smoothing to the standard errors.[18,19] Proteins with an FDR adjusted p-value < 0.05 and a fold change > 2 were considered significant.

### Tolerance tests and plasma biochemistry

Mice were fasted for 4 hrs prior to glucose (GTT) or insulin (ITT) tolerance testing. For GTT, glucose was administered intraperitoneally (i.p.) at a dose of 2 gm/kg, and for ITT, human insulin (Humulin-R) was administered i.p. at 0.6 IU/kg.[20] Following injection, blood glucose levels were recorded at 0, 30-, 60-, 90-, and 120-min using a Contour next ONE glucometer Meter (Ascensia Diabetes Care, Parsippany, NJ). Circulating total cholesterol (TC), triglycerides (TG), non-esterified fatty acids (NEFA), insulin levels, and glucagon were quantified using commercially available assay kits according to the manufacturers' instructions.

### Hematoxylin and Eosin (HE) staining

Formalin-fixed tissues were processed through graded ethanol series and xylene, followed by infiltration with liquid paraffin.[21] Paraffin-embedded tissue blocks were sectioned at 4-5 µm, and the sections were stained with hematoxylin and eosin (H&E). Images were captured in bright-field mode using an upright microscope (Leica Microsystems).

### RNA isolation and Real time PCR

Quantitative gene expression analyses for *Nlrp3* and *Gsdmd* were performed using TaqMan probes (Applied Biosystems), whereas SYBR Green chemistry was used for all other targets with primers obtained from ThermoFisher Scientific or IDT. Total RNA was extracted using TRIzol reagent (Thermo Fisher Scientific) and its quality was assessed using either a 2100 Bioanalyzer (Agilent Technologies Inc., Santa Clara CA) or a DS-11 FX nanoscale spectrophotometer (Denovix Inc.). cDNA was synthesized using High-Capacity cDNA reverse transcription kit (Applied Biosystems). Quantitative PCR was performed on a Quant Studio Real-Time PCR system (Applied Biosystems). Gene expression was normalized to 18S and calculated using the $2^{-DDCT}$ method.[22]

#### Western blot

Protein samples (10-20 μg) were run on 10% SDS-PAGE, transferred to polyvinylidene difluoride (PVDF) membranes (Thermo Fisher Scientific), blocked with 5% BSA in TBST (for 2 hrs, at RT) and then probed with primary antibodies [diluted in 1:1 ratio of TBST (pH 7.4) and 5% BSA, incubated overnight at 4 °C].[23] Membranes were later incubated with horse radish peroxidase-linked secondary IgGs (1:10,000) for 2 hrs at RT, washed and visualized using ECL detection kit (Thermo Fisher Scientific). The signals were captured along with β-actin loading control to visualize the changes in protein expression with respective controls for comparison.

#### Statistical Analysis

All data are presented as mean ± SEM (standard error of the mean). Data distributions were assessed for normality using Shapiro-Wilk and Kolmogorov-Smirnov tests. When normality assumptions were not met, non-parametric tests were applied. Accordingly, statistical comparisons were performed using Unpaired two-tailed t tests with Welch's correction (Unpaired t test), Brown-Forsythe and Welch One-Way ANOVA (One-Way ANOVA), two-way ANOVA with Sidak's multiple comparisons test (Two-Way ANOVA), Mann-Whitney test or Kruskal-Wallis test followed by uncorrected Dunn's post hoc test (Kruskal-Wallis test), as appropriate and as specified in individual figure legends. Outliers were identified using built-in GraphPad Prism routines and were excluded only when justified. A two-tailed $p < 0.05$ was considered statistically significant. All statistical analyses and graph generation were performed using GraphPad Prism (version 10.6.1).

### SUPPLEMENTAL FIGURE LEGENDS

**Figure S1. Additional metabolic parameters related to DSF effects in C57 mice.**

**(A)** Experimental outline for metabolic studies (Figures 1, 3, S2, S4, and S6).

**(B)** Body fat (%, N=12/group) at 4 weeks and **(C)** Fasting plasma glucagon in LFD, HFD, and HFDD mice (N=6/group). Kruskal-Wallis test for glucagon.

**(D)** Paired-feeding experimental design.

**(E)** Weekly BW over 15 weeks; **(F)** body fat changes at 4, 8, 12 weeks. N=4/group.

**(G, H)** GTTs during paired-feeding (8 and 14 weeks) with corresponding **(J)** AUCs. N=4.

**(I)** GTT performed after one week of ad libitum feeding (15 weeks) with corresponding **(J)** AUC. N=4.

**(K-M)** BW change (%) and GTTs after 8-week HFD feeding post BMTs in mice. N=6/group.

Data are mean ± SEM. *P<0.05, **P<0.01. N=4–12/group as indicated. A Two-Way ANOVA and One-Way ANOVA test for multiple groups, while Unpaired t test for two group comparisons was utilized unless stated. *Mann-Whitney for paired-feeding data analyses.

**Figure S2. DSF-induced weight loss in LFD-fed mice is independent of systemic metabolism.**

**(A)** Experimental outline for LFD and LFDD studies.

**(B)** Biweekly BW and **(C)** body fat (%) at 10 weeks.

**(D)** Representative H&E adipose images, adipocyte size distribution, and mean size.

**(E, F)** GTTs at 7 and 14 weeks with **(G)** corresponding AUCs.

**(H, I)** ITTs at 8 and 15 weeks with **(J)** corresponding AUCs.

**(K-M)** Fasting blood glucose, plasma insulin, and lipid profile.

**(N)** Representative pancreas H&E with islet size distribution and mean size.

Data are mean ± SEM. *P<0.05, **P<0.01. N=12/group except histological assessments where n=6/group. Statistical tests were applied wherever appropriate. Two-Way ANOVA in case of BWs and GTT-14 weeks while Unpaired t test for body fats and ITT.

**Figure S3. DSF effectively inhibited the release of intracellular protein secretome during pyroptosis in mouse BMDMs.**

**(A)** Experimental schematic: LPS priming (step 1) → nigericin (step 2) with DSF or MCC950 post-LPS. Secretome collected for proteomics. N=4/group.

**(B, C)** Venn diagrams of significantly up- and downregulated proteins.

**(D–G)** Volcano plots and pathway enrichment for indicated comparisons.

**(H–M)** DSF dose optimization: serial dilutions, outline for treatment timings, and MTT viability assays. N=4/group.

**(N)** Canonical pathway enrichment analysis (IPA).

**(O)** Immunoblot showing DSF inhibition of GSDMD cleavage (10 μM).

**Figure S4. Global deletion of GSDMD does not protect against obesity and IR.**

**(A)** Experimental design for HFD feeding in WT and *Gsdmd* knockout (KO) mice.

**(B-C)** BW progression over 16 weeks and Body fat (%) at 10 weeks.

**(D–F)** GTTs at 7 and 14 weeks with corresponding AUCs.

**(G–I)** ITTs at 8 and 15 weeks with corresponding AUCs.

**(J–L)** Fasting blood glucose, plasma insulin, and circulating lipids.

**(M)** Schematic for IDC measurements.

**(N–S)** Food and water intake, locomotor activity, and energy homeostasis parameters ($VO_2$, $VCO_2$, RER, cumulative EE, and cumulative EB) measured by IDC (n = 6–7/group).

**(T–V)** Manual food intake and BW change during diet initiation (n = 7/group).

**(W)** Immunoblot confirming loss of total and cleaved GSDMD in liver tissue (n = 3/group).

Data are mean ± SEM. *P < 0.05, *P < 0.01. n = 10 mice/group unless otherwise stated. Two-way ANOVA, Mann–Whitney test, and unpaired t test were used for GTT at 7 weeks, fasting glucose, and insulin analyses, respectively.

**Figure S5. ASO dose characterization for *Gsdmd* knockdown.**

**(A)** Study design for ASO dose optimization.

**(B–D)** *Gsdmd*, *Mcp1*, and *Nlrp3* gene expressions in pgWAT, liver, and SKM. N=3/group in case of chow mice and N=2/group for HFD mice.

**Figure S6. ASO-mediated *Gsdmd* inhibition does not improve HFD-induced metabolic dysfunction and additional parameters showing DSF-mediated metabolic protection is independent of Gasdermin D.**

**(A)** Experimental outline for CTL and *Gsdmd* ASO administration.

**(B, C)** Weekly BW and % change for 16 weeks.

**(D)** Body fat (%) at 10 weeks.

**(E, F)** GTTs at 7 and 14 weeks with **(G)** AUCs.

**(H, I)** ITTs at 8 and 15 weeks with **(J)** AUCs

**(K)** Fasting blood glucose.

**(L)** Experimental outline for IDC assessments.

**(M–Q)** IDC-assessed energy homeostasis: food/water intake, locomotor activity, $VO_2$ and $VCO_2$, RER, and cumulative EE over 48 hrs. N=8/group.

Data are mean ± SEM. *P<0.05, **P<0.01. N=11/group unless stated. Statistical tests were applied wherever appropriate. Two-Way ANOVA for % BW change, while Unpaired t test was applied for glucose and locomotor activity, respectively.

**(R)** Experimental design for DSF treatment in *Gsdmd* KO mice.

**(S-T)** GTT (7 weeks) and ITT (8 weeks) in KO mice fed with and without DSF. Data are mean ± SEM. **P<0.01. N=10/group. Two-Way ANOVA was performed here.

**Figure S7. Additional tracer and basal IDC parameters in mice fed special diets.**

**(A-D)** 13C exhalation following $13C_6$-leucine: exhalation tracer, AUC, rise and decay slopes.

**(E–J)** 12C-based metabolic parameters during $13C_6$-leucine administration: $VO_2$, $VCO_2$, RER, cumulative EB, EE, and locomotor activity. Data are mean ± SEM. *P<0.05, **P<0.01. N=6-7/group. One-Way ANOVA test was utilized for multiple groups unless otherwise stated. Unpaired t test was performed for Cum. EB between HFD and HFDD groups.

**(K-N)** Experimental outline and qPCR analyses for genes participating in lipid oxidation (*Pparα* and *Pgc-1α*) and lipogenesis (*Pparγ*) in pgWAT, liver, and SKM tissues collected between 10-12 hrs after $13C_{16}$-palmitate administration. N=5 for hepatic *Pparγ,* SKM *Pparα* and *Pparγ* expressions. Data are mean ± SEM. *P<0.05, **P<0.01. N=6-7/group and Unpaired t test was performed unless otherwise stated. Mann-Whitney test for *Pgc1α* expression in pgWAT.

**(O-T)** $VO_2$ and $VCO_2$ during cumulative, dark, and light phases, respectively in WT mice fed with different diets for ~11 weeks.

**(U–Z)** Parameters of energy homeostasis assessed during period of minimal activity: $VO_2$, $VCO_2$, RER, average EE, lipid, and carbohydrate oxidation indices in mice fed with different diets for ~11 weeks. Data are mean ± SEM. *P<0.05, **P<0.01. N=6–7/group. One-Way ANOVA test was utilized for multiple groups.
