## Supplementary material for "Disulfiram Protects Against Diet-Induced Obesity by Reprogramming Systemic Lipid Partitioning Independent of GSDMD": Reagents and Resources

### KEY RESOURCES TABLE

| REAGENT or RESOURCE | SOURCE | IDENTIFIER |
|---|---|---|
| **Chemicals, Peptides and Recombinant Proteins** | | |
| LPS from E. coli | Enzo Life Sciences | ALX-581-012 |
| Nigericin Sodium Salt | Sigma | N7143 |
| 3-(4,5-dimethylthiazol-2-yl)-2,5-diphenyltetrazolium bromide (MTT) | Thermo Fisher | M6494 |
| Pentafluorobenzyl bromide | Sigma | 90257 |
| Diisopropylethylamine | Sigma | 496219 |
| U-$^{13}C_{16}$ Palmitic acid | Cambridge Isotope Labs | CLM-409-PK |
| U-$^{13}C_6$ Glucose | Cambridge Isotope Labs | CDLM-3813-PK |
| U-$^{13}C_6$ Leucine | Cambridge Isotope Labs | CLM-2262-H-PK |
| D-Glucose | Sigma | G8720 |
| Humulin-R | El Lilly | HI-210 |
| Trizol | Thermo Fisher Scientific | 15596018 |
| 2ml bead mill tubes | Thermo Fisher Scientific | 15-340-153 |
| 13x100 mm borosilicate glass tubes | Thermo Fisher Scientific | 14-961-27 |
| **Critical Commercial Assays** | | |
| High-Capacity cDNA reverse transcription kit | ABI Biosystems | 4368813 |
| TaqMan Fast Advanced Master mix | Thermo Fisher Scientific | 4444963 |
| Fast SYBR Green Master Mix | Thermo Fisher Scientific | 4385612 |
| Total cholesterol | Infinity | TR13421 |
| Triglycerides | Infinity | TR22421 |
| NEFA | Randox | F115 |
| Insulin | ALPCO | 80-INSMSU-E10 |
| Glucagon | Crystal Chem | 81518 |
| BCA Protein Assay kit | Thermo Fisher Scientific | 23225 |
| **Animals** | | |
| C57BL/6J | JAX Laboratories | 000664 |
| C57BL/6J-*Gsdmd*$^{em1Vnce}$/J | JAX Laboratories | 032663 |
| **Diets** | | |
| LFD | Dyets | 110700 |
| LFDD | Dyets | 103530 |
| HFD | Dyets | 101920 |

| | | |
|---|---|---|
| HFDD | Dyets | 103532 |
| **Software** | | |
| NIH ImageJ | NIH | http://imagej.net |
| **Equipment/ Instruments** | | |
| QuantStudio 3 Real-time PCR System | Applied Biosystems | https://www.thermofisher.com/order/catalog/product/A28567 |
| Varioskan LUX Multimode Microplate Reader | Thermo Fisher Scientific | https://www.thermofisher.com/us/en/home/life-science/lab-equipment/microplate-instruments/plate-readers/models/varioskan.html |
| K2 Fluorescent Viability Cell Counter | Nexcelom | https://www.nexcelom.com |
| ECLIPSE Ti2 | Nikon | https://www.microscope.healthcare.nikon.com/products/invertedmicroscopes/eclipse-ti2-series |
| Element HT5 | Heska | https://www.heska.com/product/element-ht5/ |
| Micro- Cell | MicroVet Diagnostics | https://www.microvetdiagnostics.com/Products/Hematology |
| **Oligonucleotides** | | |
| ***Antisense oligonucleotides (ASOs)*** | | |
| *Gsdmd* | IONIS Pharmaceuticals | |
| *Control* | IONIS Pharmaceuticals | |
| ***TaqMan probes*** | | |
| *Nlrp3* (Mm00840904_m1) | Applied Biosystems | 4331182 |
| *Gsdmd* (Mm00509958_m1) | Applied Biosystems | 4331182 |
| Eukaryotic *18S rRNA* Endogenous Control | Applied Biosystems | 4319413E |
| Quantitative real-time PCR primers | ThermoFisher/IDT | Please see Table S1 |

**Supplemental Table S1:** Oligonucleotide sequences for RT-qPCR, related to STAR methods

| Oligo Name | Sequence | Source (Ref Seq#, NCBI) |
|---|---|---|
| **Oligos from ThermoFisher** | | |
| *Mcp1_F* | CCTGCTGTTCACAGTTGCC | NM_011333.3 |
| *Mcp1_R* | ATTGGGATCATCTTGCTGGT | |
| *Ndst1_F* | GGCAAGGAGGGCACACGCAT | NM_008306.3 |
| *Ndst1_R* | GCCGTGCTCGACAGCGAACT | |
| *Cyp2b10_F* | GGGAACCTCTTGCAGATG | NM_009999.3 |
| *Cyp2b10_R* | CCCAGGTGCACTGTGAA | |
| *Cyp27a1_F* | GGAGGATTGCAGAACTGGAG | NM_024264.4 |
| *Cyp27a1_R* | TGCGGGACACAGTCTTTACTT | |
| *Cyp7a1_F* | GTCCGGATATTCAAGGATGCA | NM_007824.2 |
| *Cyp7a1_R* | AGCAACTAAACAACCTGCCAG TACTA | |
| *Ext1_F* | GCTCTTGTCTCGCCCTTTTGT | NM_010162.2 |
| *Ext1_R* | TGGTGCAAGCCATTCCTACC | |
| *Ext2_F* | CAGGCAACATATTGAACAGC | NM_010163.2 |
| *Ext2_R* | CAGGCAACATATTGAACAGC | |
| **Oligos from IDT** | | |
| *Hmgcr_F* | GGCCTCCATTGAGATCCG | NM_008255.2 |
| *Hmgcr_R* | CACAATAACTTCCCAGGGGT | |
| *Fas_F* | GGCATCATTGGGCACTCCTT | NM_007988.3 |
| *Fas_R* | GCTGCAAGCACAGCCTCTCT | |
| *Cd36_F* | GCGACATGATTAATGGCACA | NM_001421119.1 |
| *Cd36_R* | CCTGCAAATGTCAGAGGAAA | |
| *Pparγ_F* | GATGCACTGCCTATGAGCAC | NM_011146.4 |
| *Pparγ_R* | TCTTCCATCACGGAGAGGTC | |

| | | |
|---|---|---|
| *Pparα_F* | GTCCTCAGTGCTTCCAGAGG | NM_001113418.1 |
| *Pparα_R* | GGTCACCTACGAGTGGCATT | |
| *Fabp1_F* | TCACCATCACCTATGGACCCA | NM_017399.5 |
| *Fabp1_R* | TCCAGTTCGCACTCCTCCC | |
| *Abcg5_F* | GTGCATCTTAGGCAGCTCAG | NM_031884.2 |
| *Abcg5_R* | TTCACAAACACCTCCCCTTC | |
| *Abcg8_F* | CACTGGTCATGGCTGAGAAA | NM_001347418.1 |
| *Abcg8_R* | TCCGAGGAGAACAAGCTGTC | |
| *Abcg1_F* | ATCGAATTCAAGGACCTTTCC | NM_009593.2 |
| *Abcg1_R* | TTTCCCAGAGATCCCTTTCA | |
| *Scarb1_F* | GGCTGCTGTTTGCTGCG | NM_001424495.1 |
| *Scarb1_R* | GCTGCTTGATGAGGGAGGG | |
| *Apoe_F* | AACCGCTTCTGGGATTACCT | NM_009696.4 |
| *Apoe_R* | CAGTGCCGTCAGTTCTTGTG | |
| *Ldlr_F* | GCATCAGCTTGGACAAGGTGT | NM_001252659.1 |
| *Ldlr_R* | GGGAACAGCCACCATTGTTG | |
| *Lrp1_F* | CCTGAAGGGCTTTGTGGAT | NM_008512.2 |
| *Lrp1_R* | TAGAAGTTTCCCGTCAGCCA | |
| *Pgc1α_F* | AACCACACCCACAGGATCAGA | NM_001402988.1 |
| *Pgc1α_R* | TCTTCGCTTTATTGCTCCATGA | |
| *Nr1h3_F* | GCTCTGCTCATTGCCATCAG | NM_001177730.1 |
| *Nr1h3_R* | TGTTGCAGCCTCTCTACTTGGA | |
