## Supplementary figures and images for "Disulfiram Protects Against Diet-Induced Obesity by Reprogramming Systemic Lipid Partitioning Independent of GSDMD"

### Supplemental Figure-1

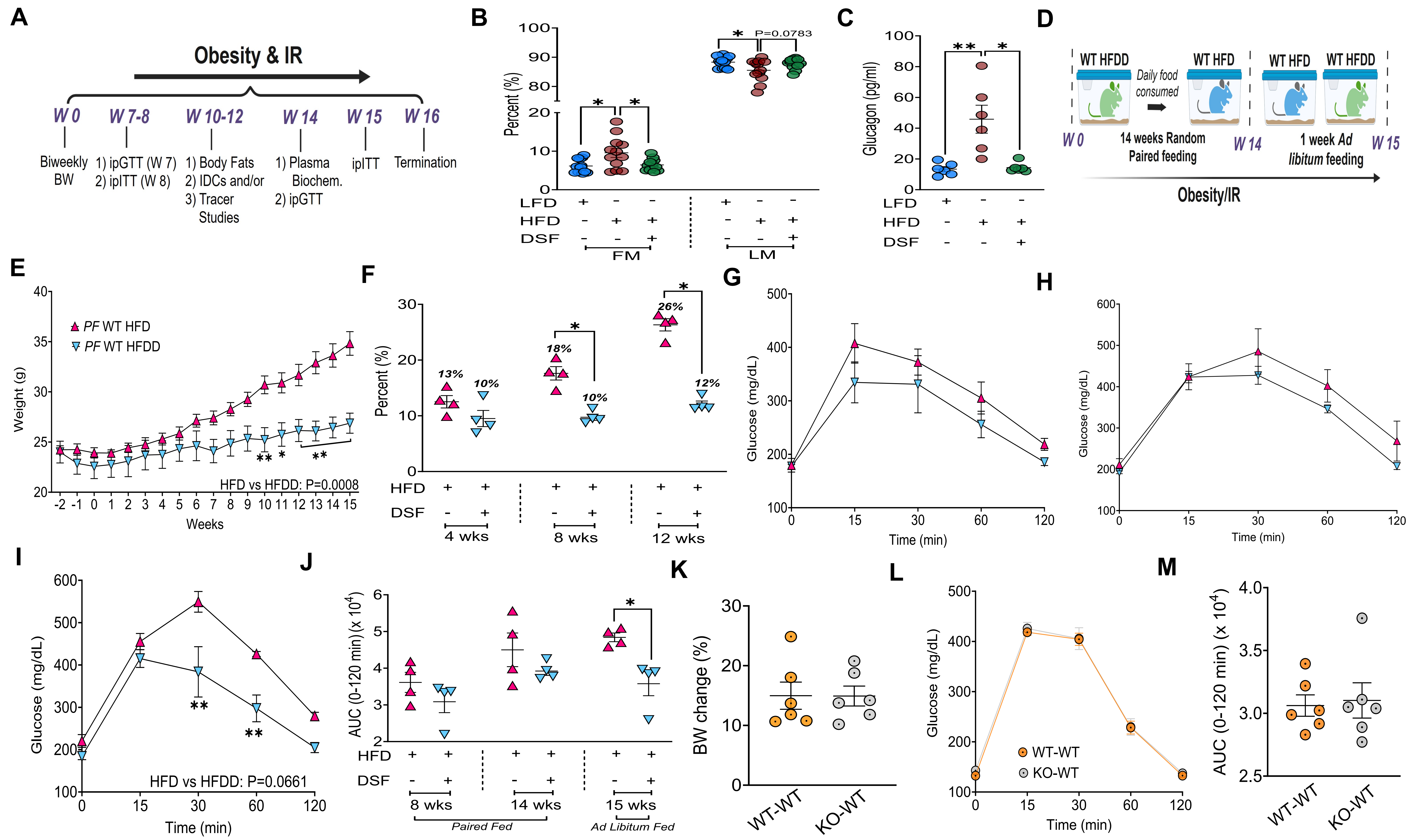

### Supplemental Figure-2

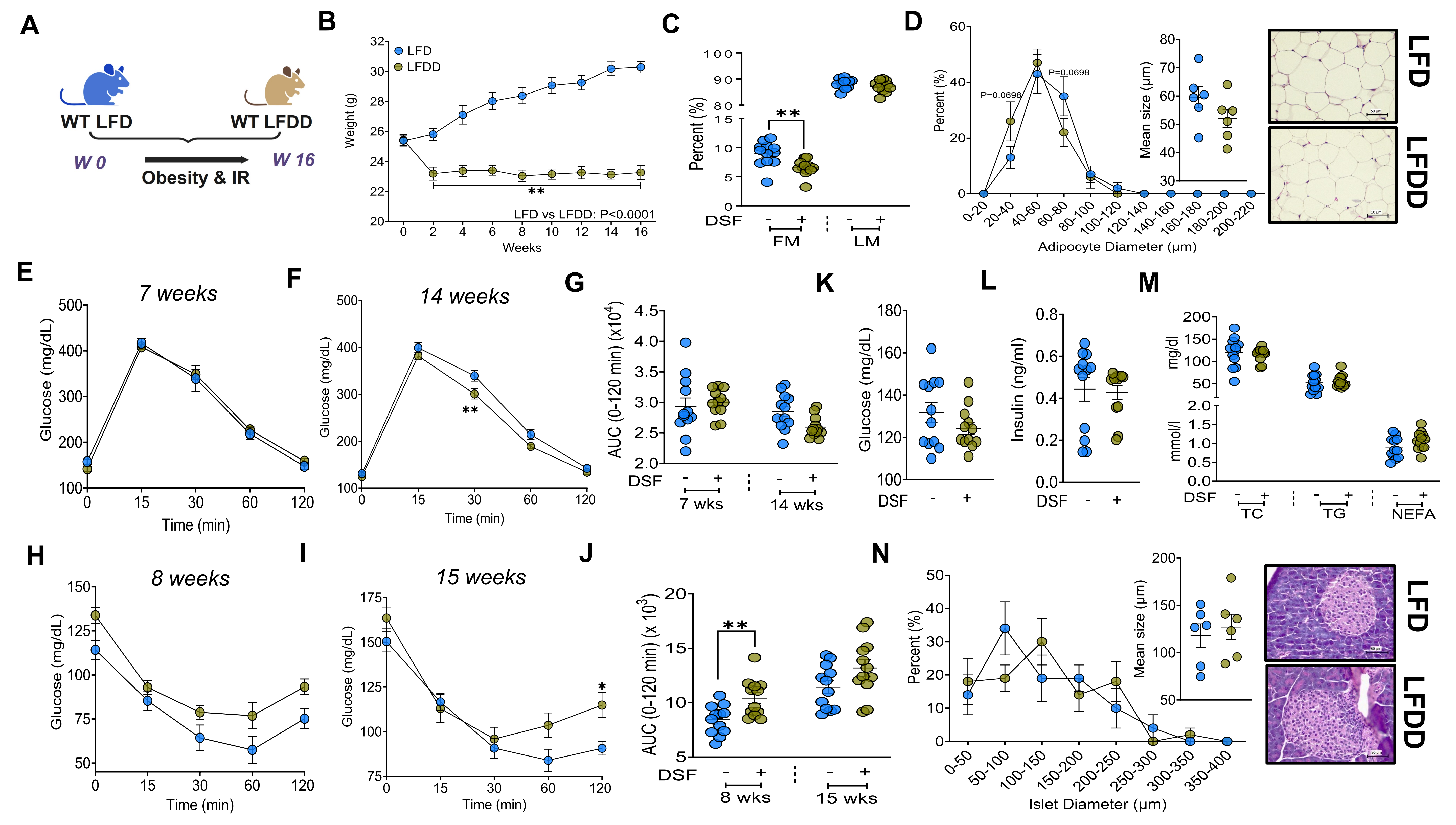

### Supplemental Figure-3

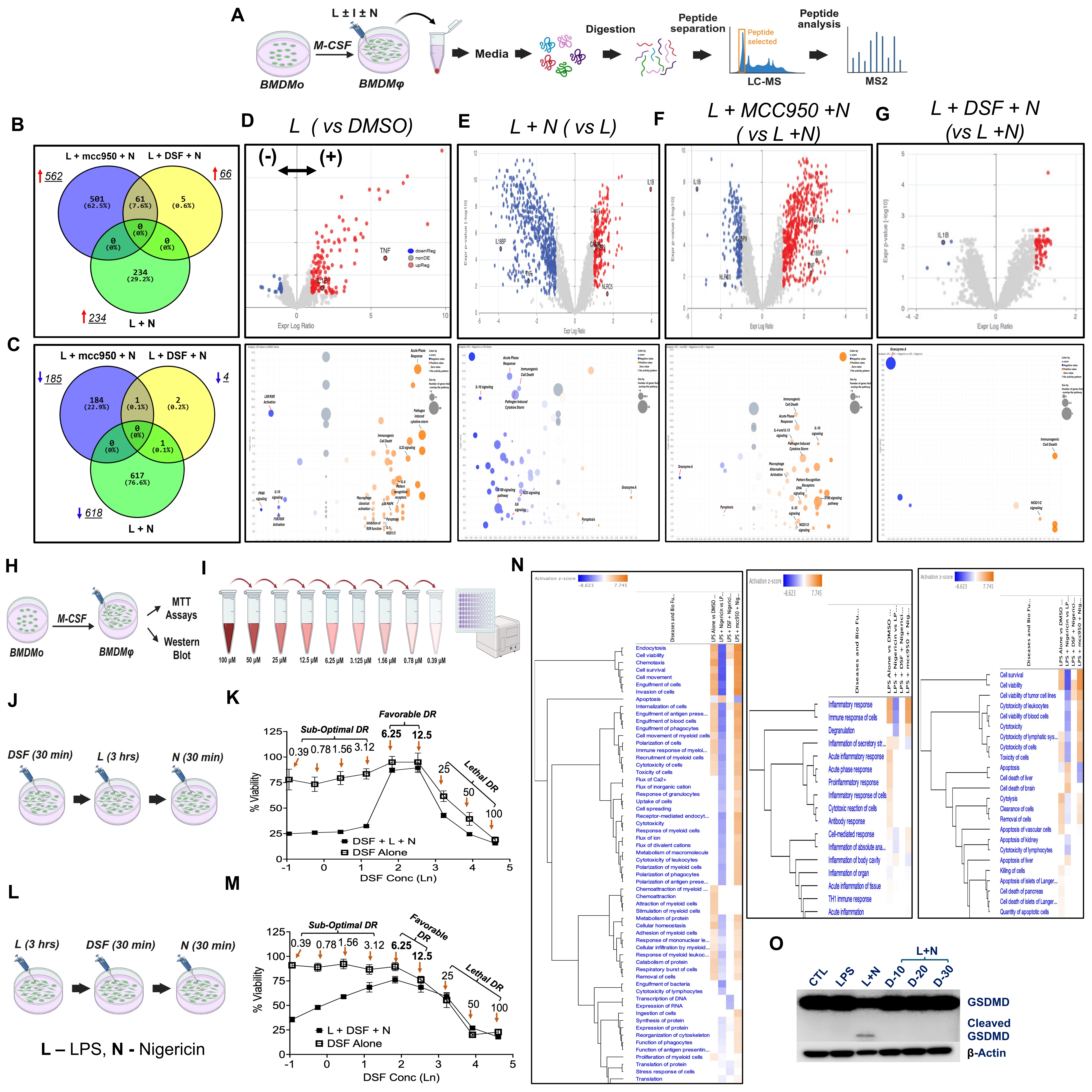

### Supplemental Figure-4

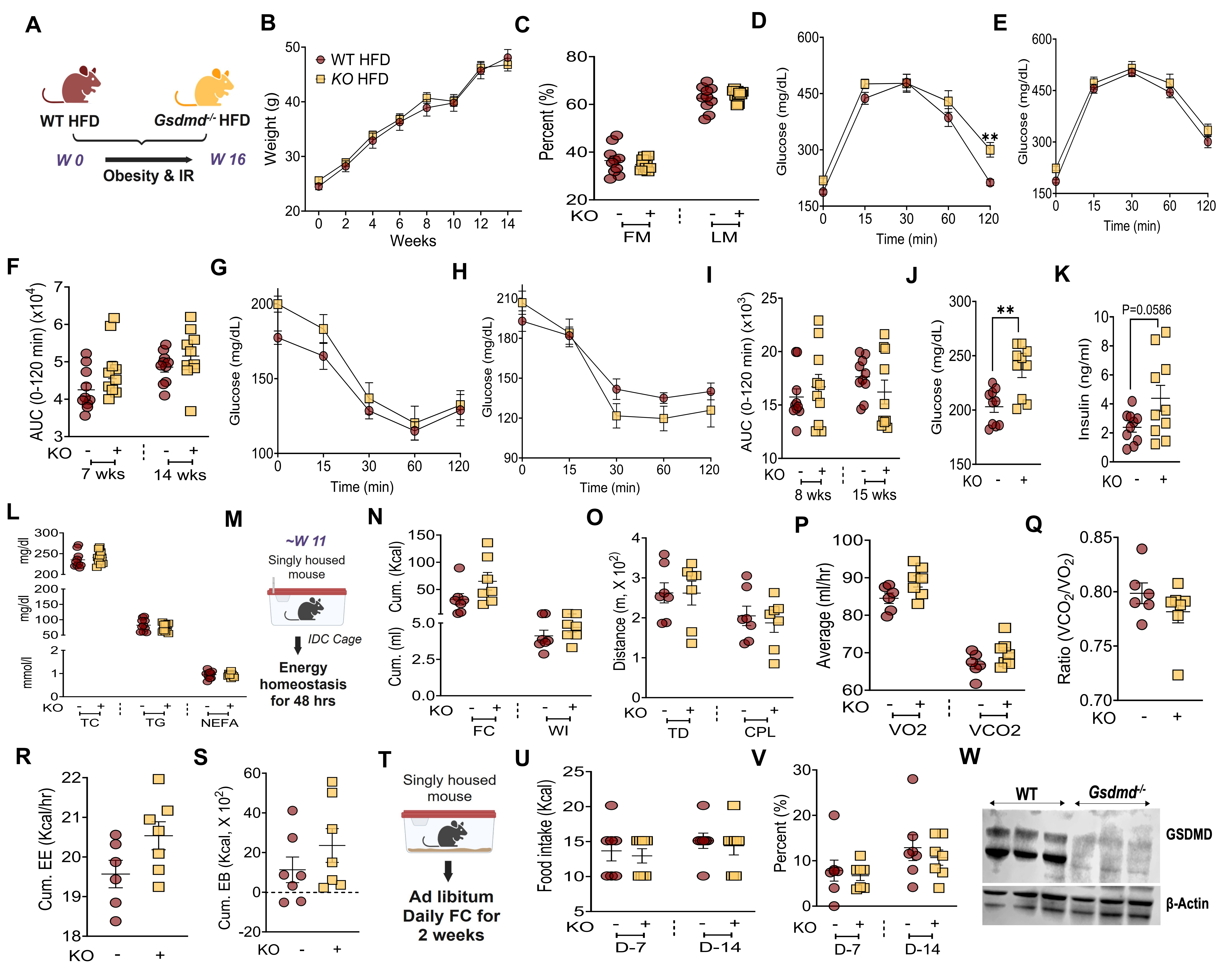

### Supplemental Figure-5

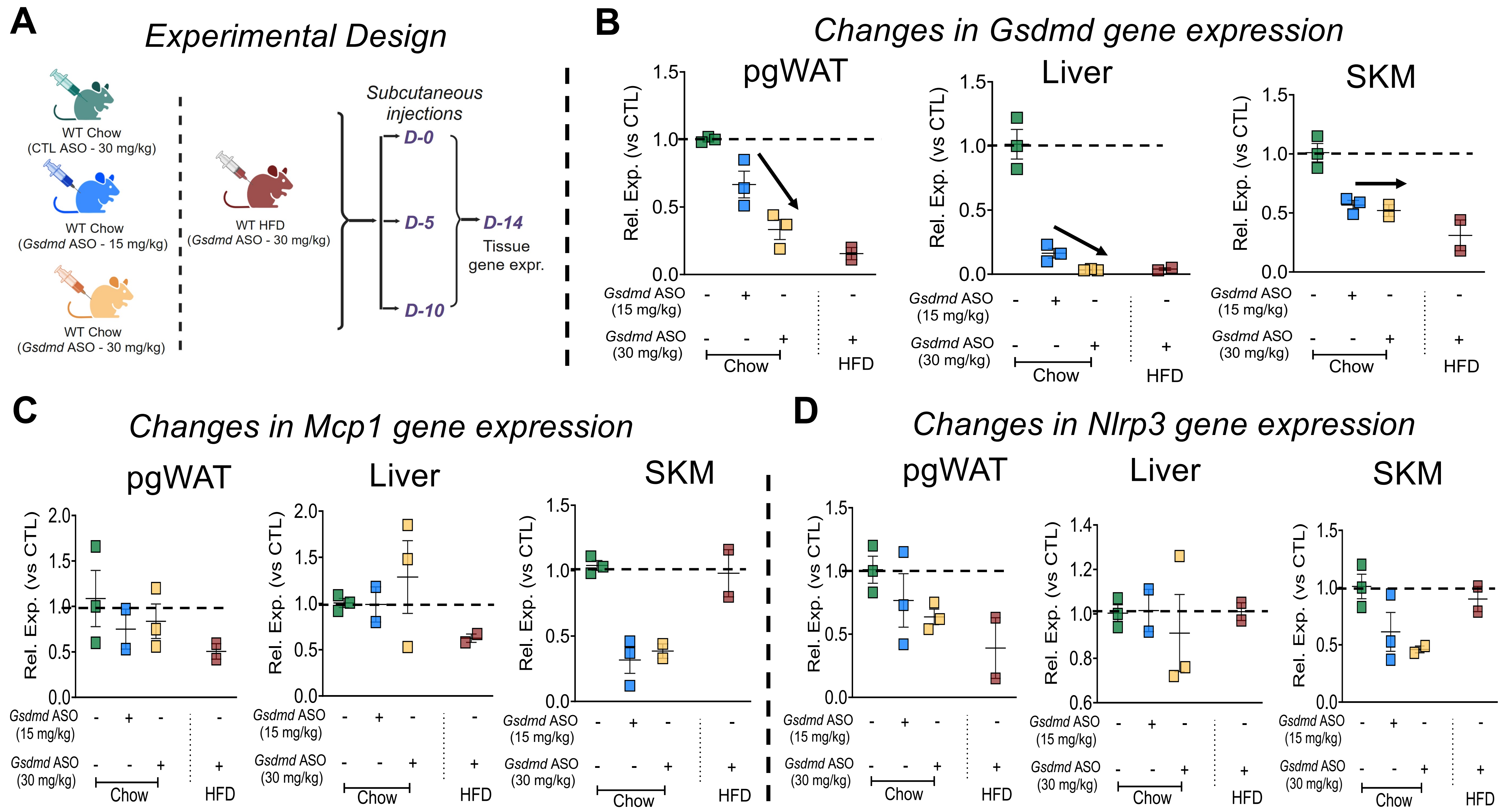

### Supplemental Figure-6

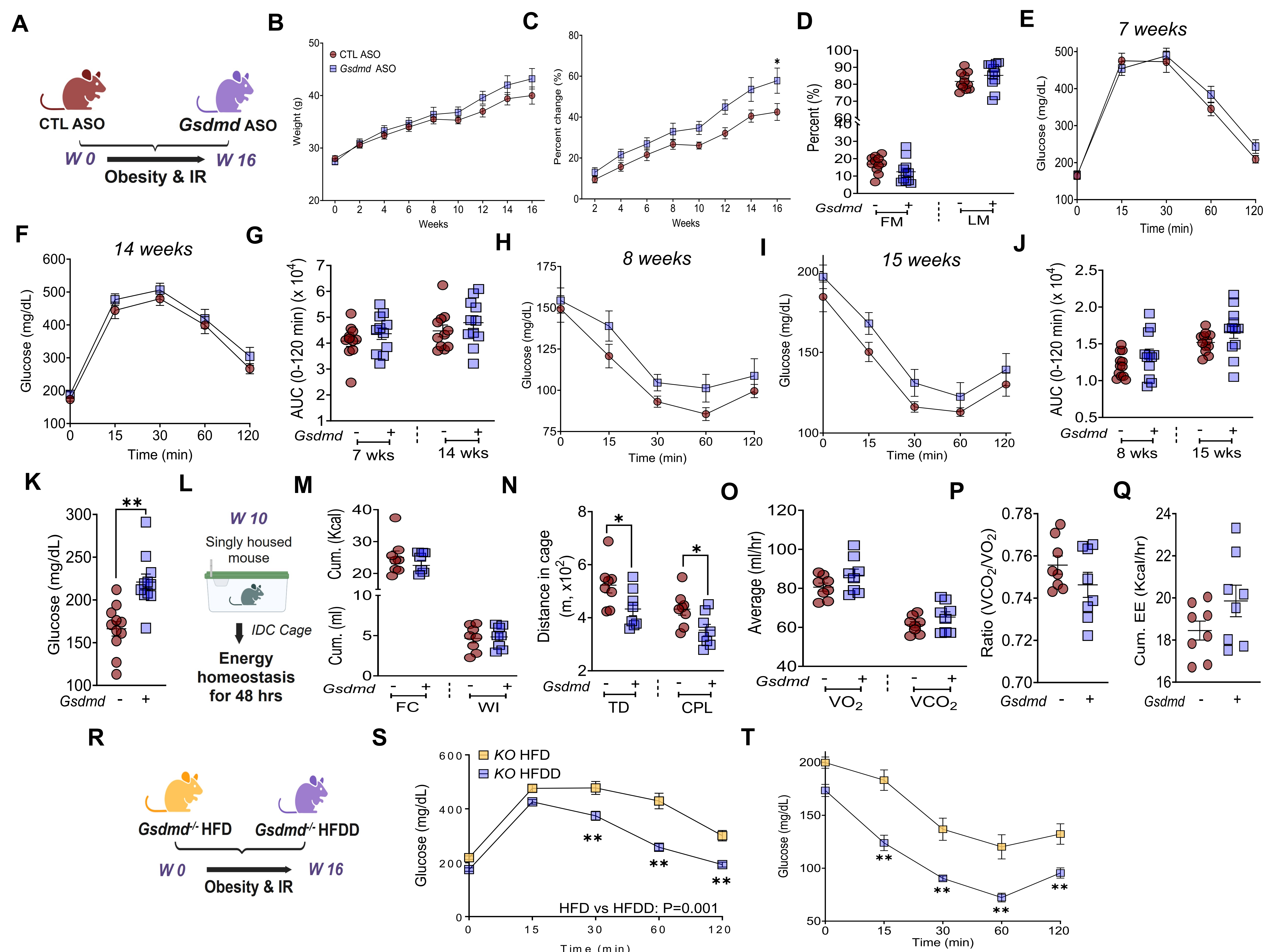

### Supplemental Figure-6

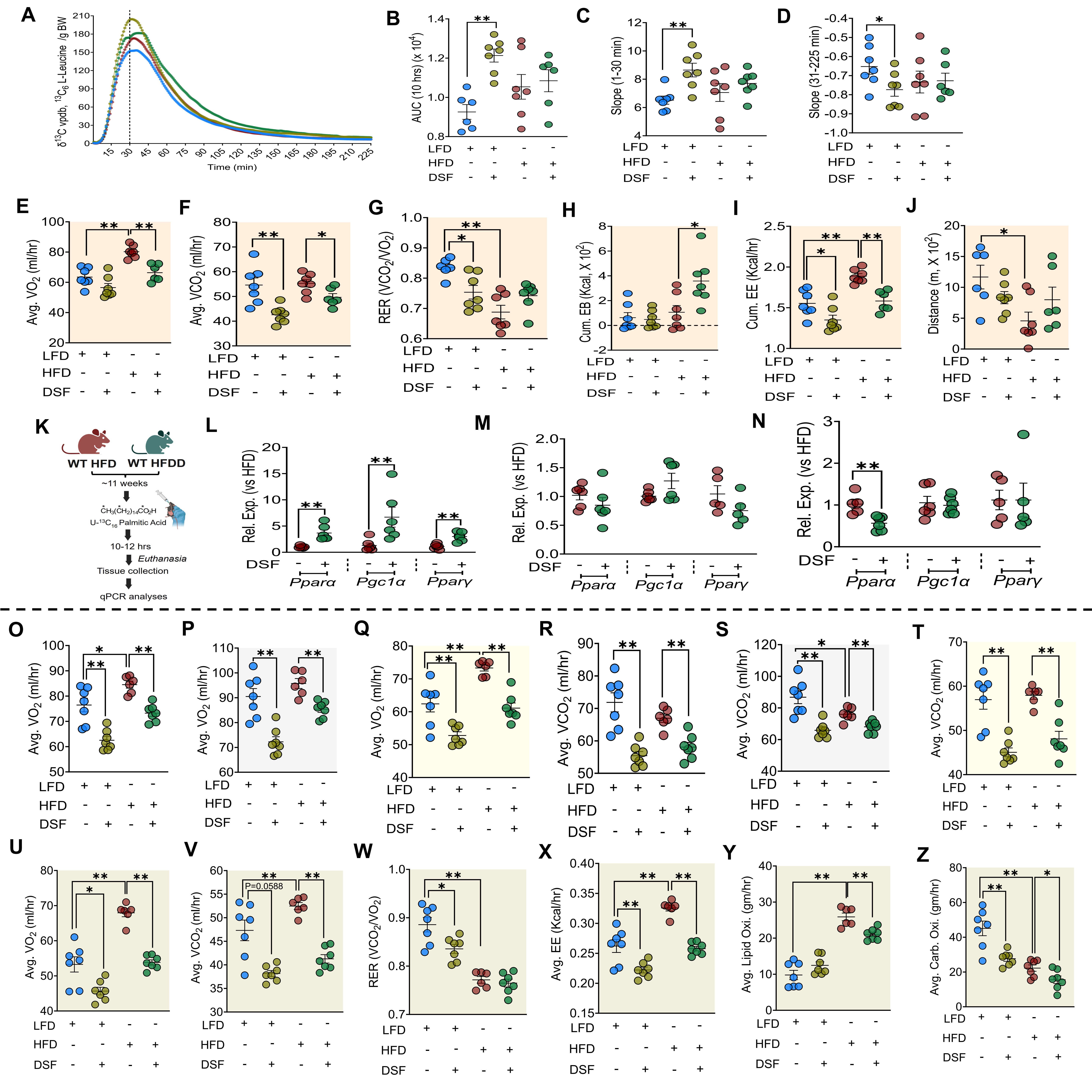
